# Serial vaccination expands and refines human CD4^+^ T cell memory

**DOI:** 10.64898/2026.04.11.716570

**Authors:** Yi-Hui Lai, Xi Su, Silina Awad, Asgar Ansari, Juneil Jang, Ali O. Saber, Huang-Wen Chen, Hannah Jung, Annesha Sarkar, Elizabeth M. Drapeau, Warren B. Bilker, Scott E. Hensley, Laura F. Su

## Abstract

CD4^+^ T cells coordinate protective immunity against pathogens. However, a major unresolved question is how human CD4^+^ T cell memory is established and evolves following primary and repeated vaccination. Using COVID-19 mRNA vaccination as a model, we tracked 50 distinct antigen-specific populations directly ex vivo with peptide–MHC class II tetramers in eight SARS-CoV-2–naïve individuals from pre-vaccine baseline through memory time points after three mRNA doses. Our findings identify the primary vaccine series as the main driver of memory pool size. It leverages pre-existing memory while preferentially recruiting high-avidity T cells, establishing an immunodominance hierarchy dominated by a small subset of precursors. Booster vaccination refines both the magnitude and quality of T cell memory. It increases select populations and enhances differentiation of subdominant CD4^+^ T cells. Populations that did not become more abundant after boosting retained their polyfunctional potential. Beyond establishing memory to the ancestral spike, vaccinations broadened responses by recruiting cross-reactive T cells recognizing viral variants. Collectively, these findings reveal how human CD4^+^ T cell memory evolves through sequential immunizations to generate a functionally diverse and broadly responsive memory repertoire against future viral challenges.

## Introduction

How vaccines and boosters confer protection remain incompletely understood. T cell-mediated immunity is a key determinant of durable protection, with CD4^+^ T cells play a key role in orchestrating coordinated immune responses. Following antigen exposure, precursor CD4^+^ T cells undergo clonal expansion and differentiate into functionally specialized subsets. Major CD4^+^ subsets include T follicular helper (Tfh) cells that provide essential signals for B cell selection and differentiation, and T helper 1 (Th1) cells, which promote antiviral effector programs and support cytotoxic CD8^+^ T cell responses ^1-3^. In the context of SARS-CoV-2 infection and vaccination, CD4^+^ T cell immunity is critical for protective immunity. Patients with milder COVID-19 infection generate higher SARS-CoV-2–specific CD4^+^ T cell responses, with earlier and more robust IFN-g induction ^4,5^. Differentiation into Th1 and Tfh effector programs inversely correlate with disease severity and are reduced among SARS-CoV-2–specific CD4^+^ T cells in hospitalized versus non-hospitalized COVID-19 patients ^6^. These studies highlight the importance of CD4^+^ T cells in protective immunity, yet key questions remain. A major limitation is the resolution of human T cell responses. Although T cells recognize discrete epitopes, antigen-specific responses are most commonly analyzed using pooled peptide stimulation ^7-11^. Data from epitope-level analyses typically focus on a few dominant responses or include limited longitudinal sampling ^12-16^. The breadth and diversity of human CD4^+^ T cell responses, and how individual populations are selectively recruited, expanded, and programed toward effector fates, remain incompletely understood.

In this study, we used COVID-19 mRNA vaccination as a model to address how CD4+ T cells memory is established and modified by primary and booster vaccines. Using peptide–MHC (pMHC) class II tetramers, we longitudinally tracked 50 distinct antigen-specific populations in eight SARS-CoV-2–naïve individuals, from the pre-vaccine baseline through memory time points following the two-dose primary series and after a booster. Our data demonstrate that primary vaccination recruits high-avidity T cells and establishes the immunodominance hierarchy in the memory pool. Dominant responses acquired a Th1 bias, whereas Tfh cells were enriched in the subdominant populations. The third vaccine fine-tunes this memory by enhancing Th1 and Tfh maturation in coordination with responsiveness to the booster. Notably, vaccination with the ancestral spike not only established memory responses to the vaccine strain, but also preferentially recruited cross-reactive T cells recognizing variant sequences. Collectively, these findings reveal how human CD4^+^ T cell memory evolves through sequential immunizations to establish a resilient memory pool capable of responding to future infections.

## Results

### Pre-existing SARS-CoV-2–specific CD4^+^ T cells include naïve and memory precursors

Eight volunteers carrying HLA alleles, DRB1*0401, *0101, and *1501, were recruited from October 2020 to April 2021 (Table S1). All individuals had no known SARS-CoV-2 infection, have not received COVID-19 vaccines, and showed a negative anti-spike antibody response at baseline (Fig. S1A). Blood samples were obtained pre-vaccination, at least 172 days after the two-dose primary series, and following the first mRNA booster (BNT162b2 or mRNA-1273) to longitudinally evaluate the establishment and subsequent modulation of memory T cells (Fig. 1A). Robust anti–spike RBD antibody responses were elicited in all donors after the primary vaccine series and were further boosted by the third mRNA vaccine dose (Fig. S1A).

**Figure 1.**
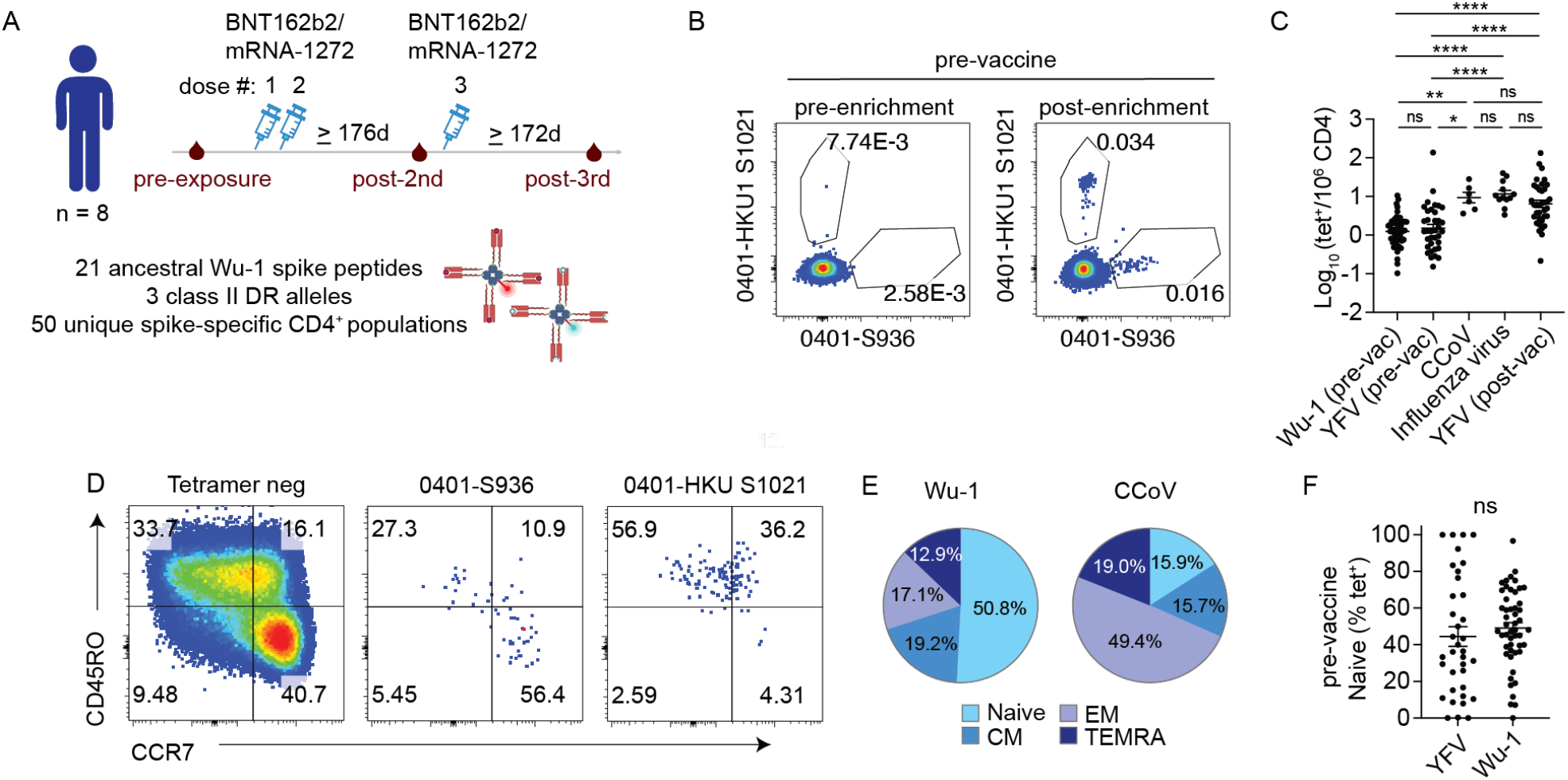
Pre-existing SARS-CoV-2–specific CD4^+^ T cells include naïve and memory precursors. (A) Overview of the study design. Vaccine-specific CD4^+^ T cells were identified in longitudinal samples from eight SARS-CoV-2–naïve donors collected before vaccination, a mean of 208 days (range 176–256) after the 2^nd^ dose, and 198 days (172–332) after the 3^rd^ dose. MHC II tetramers loaded with 21 Wuhan-1 spike peptides across three HLA-DR alleles were used to track 50 populations over time. (B) Representative direct ex vivo staining, pre- and post-magnetic enrichment of representative tetramer^+^ cells using blood obtained before vaccination. (C) The frequency of tetramer^+^ CD4^+^ T cells specific for SARS-CoV-2 Wu-1 (n = 50), YFV (n = 36), and influenza virus (n = 12) quantified by direct ex vivo staining. Vaccination status is indicated for Wu-1 and YFV. Statistical significance was determined by Kruskal–Wallis test with Dunn’s multiple-comparison correction. Data are shown as mean ± SEM. (D) Representative plots show memory subsets based on anti-CCR7 and CD45RO staining. Tetramer-negative CD4^+^ T cells are shown for comparison (left). (E) The distribution of different memory phenotype subsets among Wu-1 and CCoV-specific CD4^+^ T cells prior to vaccination: Naive (CD45RO^−^CCR7^+^); central memory (CM; CD45RO^+^CCR7^+^); effector memory (EM; CD45RO^+^CCR7^−^); and TEMRA (CD45RO^−^CCR7^−^). (F) The proportion of naïve phenotype among YFV and Wu-1 tetramer^+^ cells in pre-vaccination samples from unexposed individuals. Mann-Whitney test was performed. Data are shown as mean ± SEM. Each symbol represents the mean for a tetramer^+^ population, averaged across 1-5 independent experiments (mean = 2.5 experiments). *p < 0.05, and ****p < 0.0001.

To evaluate vaccine-specific CD4^+^ T cell responses, candidate SARS-CoV-2 spike epitopes from the ancestral Wuhan-Hu-1 (Wu-1) strain were selected based on published studies showing detectable responses in convalescent donors and based on predicted binding to HLA-DRB1*0401, *0101, and *1501 using NetMHCIIpan 4.1 from the Immune Epitope Database (IEDB) ^17,18^. To prioritize vaccine-relevant epitopes, PBMCs collected six months after vaccination were stimulated with candidate peptide pools and expanded in vitro prior to tetramer staining. Peptides yielding robust tetramer-detectable responses were validated by direct ex vivo staining using baseline and post–second dose samples. Spike-specific populations with post-vaccination frequencies exceeding the mean pre-vaccination baseline across all populations were selected for longitudinal analysis. In total, we generated tetramers for 21 peptides across three HLA-DR alleles and tracked 50 distinct antigen-specific CD4^+^ T cell populations in the blood before and after vaccination (Fig. S1B) (Table S2).

We first evaluated the Wu-1–specific precursor repertoire using pre-vaccine blood samples (Fig. 1B, Fig. S1C). For comparison with antigen-experienced T cells, we analyzed CD4^+^ T cell responses to two epitopes from the common cold coronavirus HKU1 (CCoV). To contextualize spike-specific precursors against other well-characterized virus-specific populations, we included historical datasets of yellow fever virus (YFV)–specific CD4^+^ T cells before and after YFV vaccination, as well as influenza hemagglutinin (HA_306_)–specific T cells from our prior studies ^19,20^. The comparison between pre-vaccine Wu-1 and YFV-specific CD4^+^ T cells showed similar baseline precursor frequencies, both of which were significantly lower than those observed in post-exposure CCoV, influenza virus, and YFV-specific populations (Fig. 1C). To characterize T cell differentiation, we broadly classified tetramer^+^ cells by CD45RO and CCR7 expression into naïve (CD45RO^−^CCR7^+^), central memory (CM; CD45RO^+^CCR7^+^), effector memory (EM; CD45RO^+^CCR7^−^), and TEMRA (CD45RO^−^CCR7^−^) subsets (Fig. 1D). As expected, most CCoV-specific CD4^+^ T cells expressed memory markers, with the majority expressing an EM phenotype (Fig. 1E). Spike tetramer^+^ pre-vaccine precursors contain a subset of memory phenotype T cells, in agreement with prior studies from us and others ^20-28^. The frequency of pre-existing memory cells within a tetramer^+^ population positively correlated with total population frequency, supporting a history of antigen-driven expansion and differentiation (Fig. S1D). Despite differences in exposure history, which includes closely related CCoV for SARS-CoV-2 but not to related viruses for YFV, Wu-1 spike and YFV-specific CD4^+^ T cell precursors exhibited similar proportions of naïve cells (Fig. 1F). These findings indicate a heterogeneous baseline spike precursor repertoire and suggest that repeated exposure to antigenically related pathogens does not substantially deplete the naïve T cell pool.

### Primary COVID-19 vaccines recruit high avidity CD4^+^ T cells

To examine primary memory responses, we performed direct ex vivo tetramer staining to compare Wu-1 spike-specific CD4^+^ T cells in pre-vaccination samples with those collected 6-8 months after the initial two-doses of COVID-19 mRNA vaccines (Fig. 2A). Vaccination induced a significant increase in Wu-1 spike-specific CD4^+^ T cells (Fig. 2B). The efficiency by which each population established a memory pool, quantified as the fold-change (FC) in frequency before and after vaccination (post-2^nd^/pre), strongly correlated its post-vaccine population size as frequency per million CD4^+^ T cells (Fig. 2C). On average, COVID-19 vaccination induced 148-fold increase in Wu-1 spike-specific CD4^+^ T cells, with individual populations increasing from 2 to 1092-fold from baseline. In contrast, minimal changes were observed for T cells recognizing the selected CCoV epitopes (Fig. 2D). In prior studies on YFV vaccine, we observed limited gain in populations dominated by pre-existing memory T cells after vaccination ^19^. Comparative analyses between YFV and SARS-CoV-2 showed that the negative correlation between fold expansion and pre-existing memory observed in YFV responses was absent for Wu-1 spike responses after vaccination (Fig. S2). The modest correlation between the frequencies of pre-existing memory and post-vaccine T cells suggest a positive contribution of the baseline antigen-experienced cells to the subsequent vaccine response (Fig. 2E). In addition, vaccination reorganized the immunodominance hierarchy of the baseline repertoire. The ten most expanded post-vaccine populations arose from fewer than 18% of precursor cells yet accounted for over 70% of the total primary memory pool (Fig. 2F). Using median staining intensity (MFI) of tetramers to approximate the strength of TCR-ligand interaction, we showed a significant increase in average tetramer MFI after vaccination (Fig. 2G). The gain in tetramer MFI correlated with the FC in tetramer frequency, consistent with preferential recruitment of higher avidity T cells (Fig. 2H). Together, these data suggest that mRNA COVID-19 vaccines build on pre-existing antigen-experienced T cells and selectively expand high-avidity clones to establish a robust memory pool that redefines the immunodominance hierarchy.

**Figure 2.**
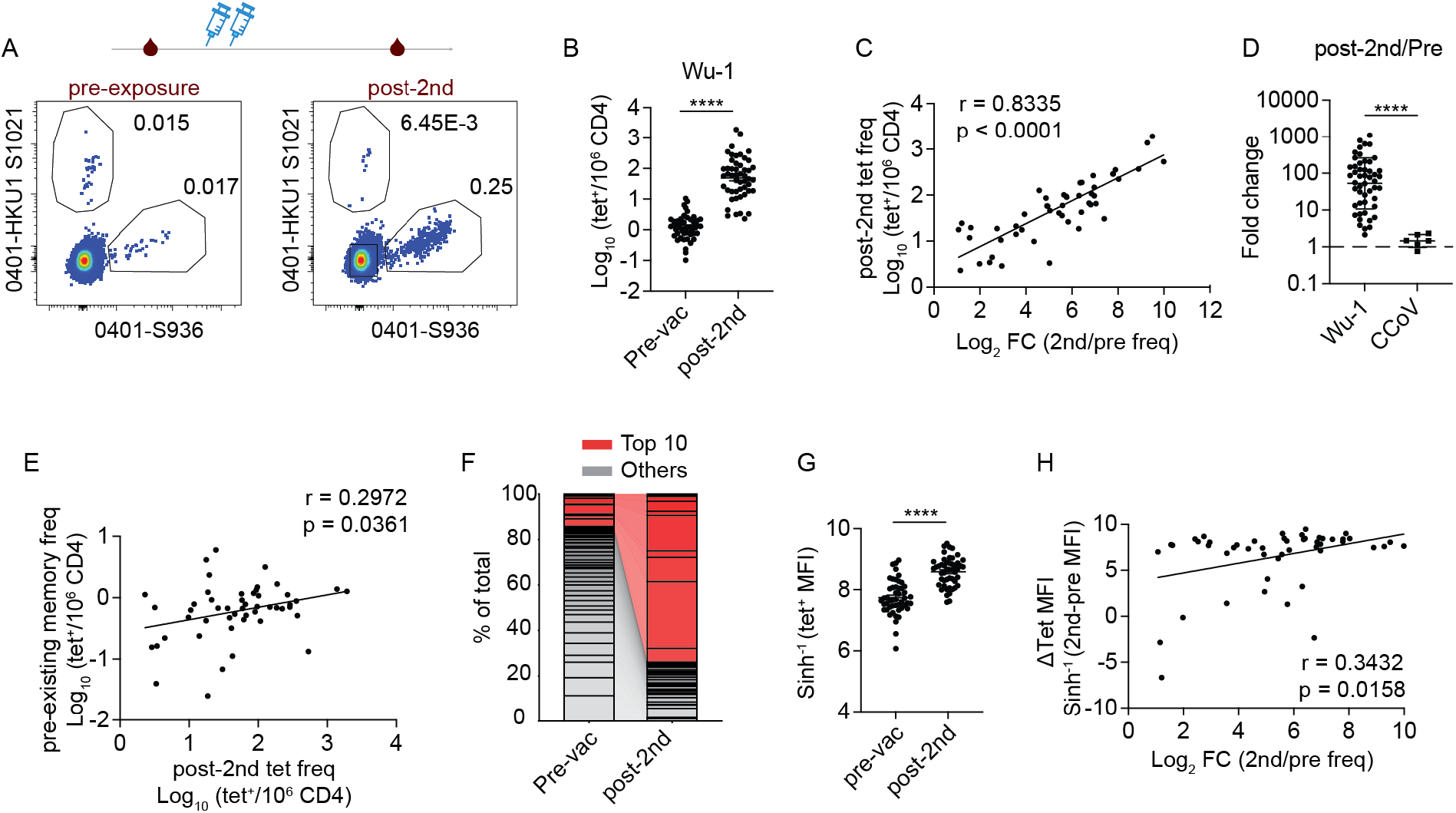
Primary COVID-19 vaccines recruit high avidity CD4^+^ T cells. (A) Representative direct ex vivo tetramer staining of Wu-1- and HKU-1–specific CD4^+^ tetramer^+^ T cells at the indicated time points. (B) Frequency of Wu-1 tetramer^+^ CD4^+^ T cells pre-vaccination and after the 2^nd^ vaccine dose (post-2^nd^). Paired t test was performed. Data are shown as mean ± SEM. (C) Correlation between the gain in frequency as a fold change from the pre-vaccine baseline and tetramer^+^ T cell frequency after the 2^nd^ dose (n= 50). Pearson correlation was performed. (D) Plot shows fold change in frequency of Wu-1 and CCoV^+^ tetramer^+^ T cells relative to pre-vaccine baseline. Mann-Whitney test was performed. Data are shown as geometric mean ± geometric SD. (E) Correlation of post-2^nd^ dose tetramer^+^ frequency with its baseline pre-existing memory frequency. Spearman correlation was used. (F) Relative abundance of each SARS-CoV-2 Wu-1 population as a percentage of total tetramer^+^ cells at each time point. The top 10 populations expanded after the second dose are highlighted in red and linked to their baseline counterparts. (G) Plot summarizes Wu-1 tetramer MFI before and after the 2^nd^ vaccine dose. Paired t-test was performed. Data are shown as mean ± SEM. (H) Correlation between fold change in tetramer frequency and the change in Wu-1 tetramer MFI before and after the 2^nd^ vaccine dose. Spearman correlation was used. Each symbol represents the mean for a tetramer+ population, averaged across 1-5 independent experiments. ****p < 0.0001.

### Vaccine responsiveness shapes T cell differentiation states after primary COVID-19 vaccinations

Stratifying tetramer^+^ populations by into top and bottom thirds by fold-change identified dominant and subdominant subsets with significantly different post-vaccination frequencies (Fig. 3A-B). Dominant populations were enriched for EM and TEMRA phenotypes, whereas subdominant responses retained more naïve-like and central memory features (Fig. 3C, S3A-D). To examine other differentiation states, cells were stained for markers of Th1 (CXCR3) and Tfh cells (CXCR5 and PD-1) (Fig. 3D). We showed that primary vaccine series increased both the percentage and frequency of tetramer^+^ Th1 and circulating Tfh (cTfh) polarized subsets (Fig. 3E-F). Notably, the magnitude of response was associated with distinct phenotypic biases. Populations with greater fold expansion were Th1-skewed and contained proportionally fewer cTfh cells, whereas limited expansion was linked to enrichment of the CXCR5^+^PD-1^+^ subset (Fig. 3G-H). When divided by tertiles, immunodominant populations within the upper third of responses contained a significantly higher frequency of Wu-1-specific CXCR3^+^ Th1-like T cells. These abundant specificities also contributed to more cTfh memory cells overall, even though a smaller fraction expressed the CXCR5^+^PD-1^+^ phenotype compared with subdominant populations (Fig. 3I-J). These data demonstrate that primary COVID-19 mRNA vaccination generates qualitatively distinct memory CD4^+^ T cells that vary with the magnitude of the response.

**Figure 3.**
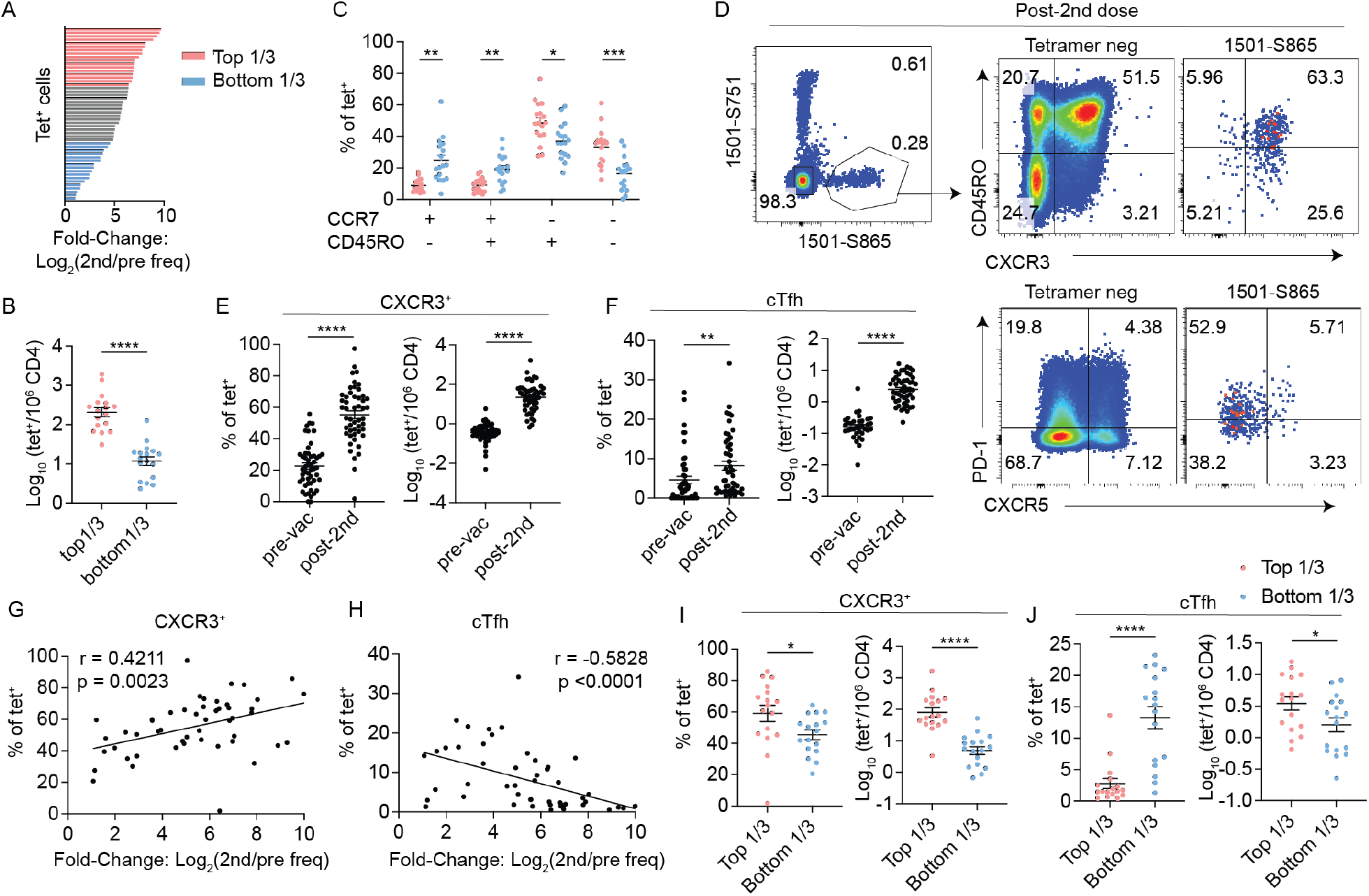
Vaccine responsiveness shapes CD4^+^ T cell memory differentiation. (A) Wu-1 tetramer^+^ CD4^+^ T cell populations ranked by fold change from baseline after the 2^nd^ dose. Top third in pink, bottom third in blue. (B) Plot comparing tetramer^+^ CD4^+^ T cell frequencies at the post-2^nd^ time point for the top 1/3 versus bottom 1/3 populations (n = 17 in each group). Welch’s t-test was performed. (C) Distribution of memory phenotype among top 1/3 and bottom 1/3 tetramer^+^ populations at the post-2^nd^ vaccine time point. RM two-way ANOVA with Sidak’s multiple-comparison test were performed. (D) Representative plots show gating for Th1-like (CXCR3^+^) and cTfh-like (CXCR5^+^PD-1^+^) phenotypes post 2^nd^ vaccine dose. (E) Frequency and proportion of CXCR3^+^ cells among Wu-1 tetramer^+^ CD4^+^ populations pre-vaccination and after the 2^nd^ vaccine dose (n = 50). Paired t-test was used for percentage and Wilcoxon test for frequency, based on Shapiro-Wilk normality testing. (F) Plots summarize cTfh frequency and percentage before and vaccination. Wilcoxon test was used. (G) Correlation between FC from baseline and the proportion of CXCR3^+^ cells after the 2-dose vaccination. Pearson correlation was used. (H) Correlation between FC and cTfh percentage post-vaccination. Spearman correlation was used. (I) Plots compare CXCR3^+^ proportions (left) and frequencies (right) in the top and bottom tertile post-2^nd^ dose. Welch’s t-test was performed. (J) Plots compare cTfh proportions and frequency after vaccination as in (I). Mann-Whitney test was used for percentage and Welch’s t-test for frequency, based on Shapiro-Wilk normality testing. Each symbol represents the mean for a tetramer+ population, averaged across 1-5 independent experiments. Data are represented as mean ± SEM. *p < 0.05, ** p< 0.01, ***p < 0.001, and ****p < 0.0001.

### Immunodominant hierarchy of primary memory is preserved following booster vaccine

The effects of booster vaccine on T cell response remain unresolved. To address this, we analyzed samples collected after a third COVID-19 mRNA vaccine dose to examine the booster effect (Fig. 4A). Booster immunization resulted in further increase in spike-specific CD4^+^ T cells (Fig. 4B). However, the magnitude of this increase was, on average, 85-fold smaller than that observed after the first two doses (Fig. 4C). Primary COVID-19 vaccination induced an average 148-fold increase from the pre-vaccine baseline (2nd/pre, range 2 to 1,094), whereas the booster expanded the antigen-experienced repertoire by at most 6.5-fold relative to pre-booster frequencies (3rd/2nd). Despite this modest overall numerical gain, greater fold-change after boosting correlated with higher post-booster frequencies (Fig. 4D). When tetramer^+^ T cells were classified by booster-induced fold change as expanded (FC > 2) or stable/contracted (FC < 1), expanded populations were significantly more abundant after the booster (Fig. 4E-F). The magnitude of response to the first two doses did not predict responsiveness to the booster (Fig. S4A). Unlike earlier responses, tetramer MFI did not change significantly after the third vaccine dose, suggesting that high-avidity populations were primarily selected during the initial series (Fig. 4G). Consistent with this, the immunodominance hierarchy was relatively stable, with tetramer frequencies after the primary series strongly correlating with those after the third dose (Fig. 4H). Rather than driving major changes, the booster fine tunes the memory repertoire. We observed that populations within the FC > 2 group generally remained in or moved into the highest-frequency tier after the booster, whereas FC < 1 populations were less abundant and either stayed in or shifted to the bottom third (Fig. S4B). The pattern of immunodominance was frequently shared between donors after vaccinations. For populations recognizing the same pMHC in at least two donors, responses to the same pMHC from different individuals preferentially grouped into the same tertile after the second and third vaccines doses, but not in the pre-vaccine baseline (Fig. S4C). During the study, six of eight volunteers were infected with COVID-19 between the second and third doses. Grouping post-booster tetramer^+^ frequencies by prior infection status revealed similar levels in both groups, indicating that infection did not substantially affect the magnitude of memory generated by the booster (Fig. S4C). Collectively, these data indicate that immunodominance hierarchy is established early and shared between individuals. The third COVID-19 mRNA dose refines the memory repertoire to provide a modest increase in the overall spike-specific responses.

**Figure 4.**
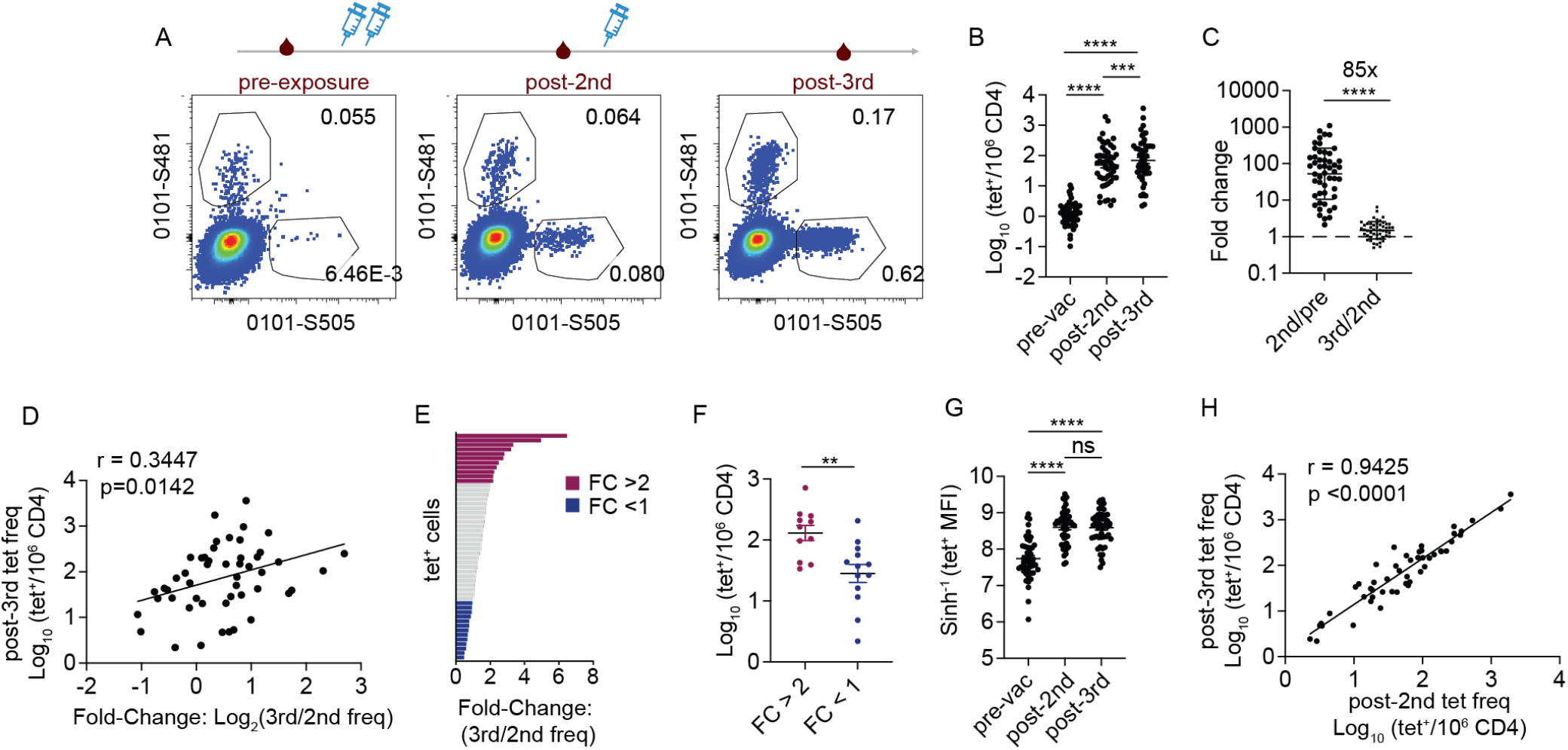
Booster vaccination increases T cell frequency while preserving the established immunodominance hierarchy. (A) Representative plots showing Wu-1 tetramer^+^ staining across the indicated time points. (B) Frequency of Wu-1 tetramer^+^ CD4^+^ T cells at indicated time points (n=50). RM one-way ANOVA with Tukey’s multiple comparison test was performed. (C) Fold change in tetramer^+^ cell frequency comparing response to the primary vaccine series (2^nd^/pre) to the booster response (3^rd^/2^nd^). Mann-Whitney test was performed. Geometric mean ± geometric SD. (D) Correlation between responsiveness to the booster as a fold change from post-2^nd^ dose baseline and tetramer^+^ frequency after the booster. Pearson correlation was performed. (E) Distribution of Wu-1 tetramer^+^ populations ranked by fold change to the booster from the post-2^nd^ dose baseline. (F) Comparison of tetramer^+^ population frequencies after the 3^rd^ vaccine dose, grouped by FC in response to the booster, FC > 2 (n = 11) and FC <1 (n = 13). Welch’s t-test was performed. (G) Tetramer MFI of Wu-1 tetramer^+^ populations at pre-vaccination and following the 2^nd^ and 3^rd^ doses of the COVID-19 vaccines. RM one-way ANOVA with Tukey’s multiple comparison test was performed. (H) Correlation between tetramer^+^ T cell frequencies after the 2^nd^ and 3^rd^ vaccine doses. Association was measured by Pearson correlation. Each symbol represents the mean for a tetramer^+^ population, averaged across 1-5 independent experiments. Data are represented as mean ± SEM. *p < 0.05, ** p< 0.01, ***p < 0.001, and ****p < 0.0001.

### COVID booster selectively augments Th1-like and Tfh-biased differentiation

Next, we examined how booster vaccination modifies T cell differentiation (Fig. 5A). Following the primary COVID-19 vaccination series, vaccine-specific CD4^+^ T cells predominantly exhibited an effector memory (EM) phenotype, which was further increased by the booster and accompanied by a relative reduction in naïve-like and TEMRA subsets (Fig. 5B-C). With respect to cell fate polarization, CXCR3, CXCR5, and PD-1 expression measured across successive time points showed an increase in Wu-specific cTfh cells and stable CXCR3^+^ T cell frequency, with an overall increase in cTfh-to-Th1 ratio after boosting (Fig. 5D-H). Notably, booster-induced changes appear to selectively impact subdominant populations that showed weaker responses to the primary vaccine series. We examined phenotypic differences between populations in the top and bottom thirds based on their response to primary vaccine series. The third vaccine dose did not further increase CXCR3^+^ or cTfh frequencies in the top third of populations with the strongest fold-change responses, but it enhances these subsets among the less-expanded populations in the bottom third (Fig. 5J). In contrast, EM differentiation increased similarly across groups (Fig. 5K), suggest that boosting preferentially modulates CD4^+^ T cell polarization rather than broadly altering effector memory differentiation.

**Figure 5.**
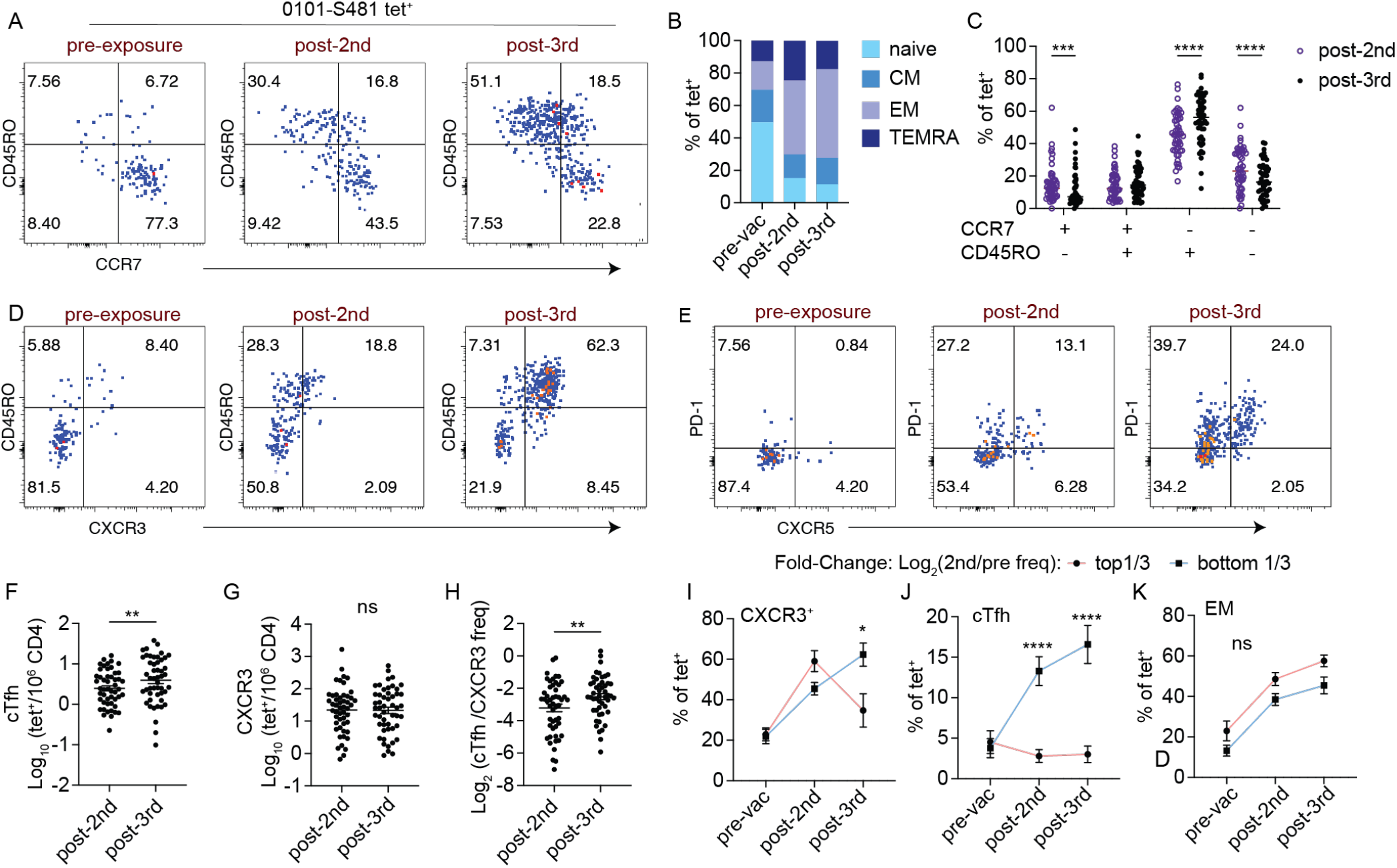
Booster vaccination augments further memory differentiation. (A) Plots show longitudinal change in CD45RO and CCR7 staining on a representative Wu-1 tetramer^+^ population pre-vaccination, post-two-dose series, and after booster. (B) Relative percentage of naïve, CM, EM, and TEMRA phenotypic subsets among Wu-1 tetramer^+^ populations across time points. (C) Plot compares the percentages of tetramer^+^ cells with the indicated CCR7 and CD45RO expression following the 2^nd^ and the 3^rd^ vaccine doses. RM two-way ANOVA with Sidak’s multiple-comparison test were used. (D-E) Representative plots of 0101-S481 tetramer^+^ CD4^+^ T cells showing CD45RO and CXCR3 (D) or CXCR5 and PD-1 (E) expression at each time point. (F) The frequencies of CXCR5^+^PD-1^+^ tetramer^+^ cells after the 2^nd^ and 3^rd^ vaccine doses. Paired t-test was performed. (G) The frequencies of CXCR3^+^ tetramer^+^ cells after the 2^nd^ and 3^rd^ vaccine doses. Wilcoxon test was used. (F) The ratio of cTfh to CXCR3^+^ T cell frequencies after the 2^nd^ and 3^rd^ vaccine doses. Paired t-test was performed. (I-K) The percentages of CXCR3^+^ (I), cTfh (J), and EM (K) subsets within tetramer^+^ populations across time points, grouped by fold-change response to the first two doses. RM two-way ANOVA with Sidak’s multiple-comparison test were used to compare top and bottom 1/3 at each time point. Each symbol represents the mean for a tetramer^+^ population, averaged across 1-5 independent experiments. Data are represented as mean ± SEM. *p < 0.05, ** p< 0.01, ***p < 0.001, and ****p < 0.0001.

### Response to booster is associated with distinct functional states

To determine if phenotypic maturation were related to responsiveness to booster, we examined the relationship between Th1-like and Tfh phentypes and the magnitude of booster response. We showed that the FC in tetramer frequency from the pre-booster baseline positively correlated with the frequency of CXCR3^+^ and CXCR5^+^PD-1 subsets within tetramer^+^ populations after the 3^rd^ dose (Fig. 6A-B). Stratifying by FC expansion also showed an enrichment of Th1-like and cTfh cells within booster-responsive group (FC > 2) compared with the nonresponsive group (FC < 1) (Fig. 6C-D). To evaluate functional potential, post-booster PBMCs were stimulated with PMA and ionomycin, followed by tetramer and antibody staining. 15,432 tetramer labeled T cells were identified by manual gating and combined for analyses using the Spectre pipeline workflow ^29^. As expected for antigen-experienced T cells, memory markers, CD45RO and CD95, were broadly expressed (Fig. S5A-C). Rare cells lacking both CD45RO and CD95 expression could also be detected, suggesting preservation of naïve-like states in a minority of tetramer^+^ T cells (clusters 4 and 10). Cytokines, IFN-γ, TNF-α, and IL-2, were expressed by cells occupying discrete yet partially overlapping regions of the UMAP. Cells in clusters 2 and 3 expressed Granzyme A, suggesting cytotoxic potential. Notably, populations in FC > 2 and FC < 1 groups showed distinct spatial distributions (Fig. 6E). FC > 2 cells localized to UMAP regions enriched for IFN-γ staining, whereas FC < 1 cells visually mapped to distinct regions with high IL-2 and TNF-α expression (Fig. 6F). Manual gating confirmed increased IL-2 expression in unexpanded populations, with enrichment of IFN-γ^−^IL-2^+^TNF-α^+^ and IFN-γ^+^IL-2^+^TNF-α^+^ subsets within these IL-2^+^ cells (Fig. 6G–H). The frequency of IL-2^+^TNF-α^+^IFN-γ^+^ cells inversely correlated with fold expansion after the third dose, suggesting that polyfunctionality is preferentially retained in less-responsive populations (Fig. 6I). These data reveal distinct functional states associated with booster responses. The booster fine-tunes the functional phenotype of vaccine-specific memory, shifting cells from IL-2 dominant polyfunctional state toward defined Th1-like and cTfh differentiation.

**Figure 6.**
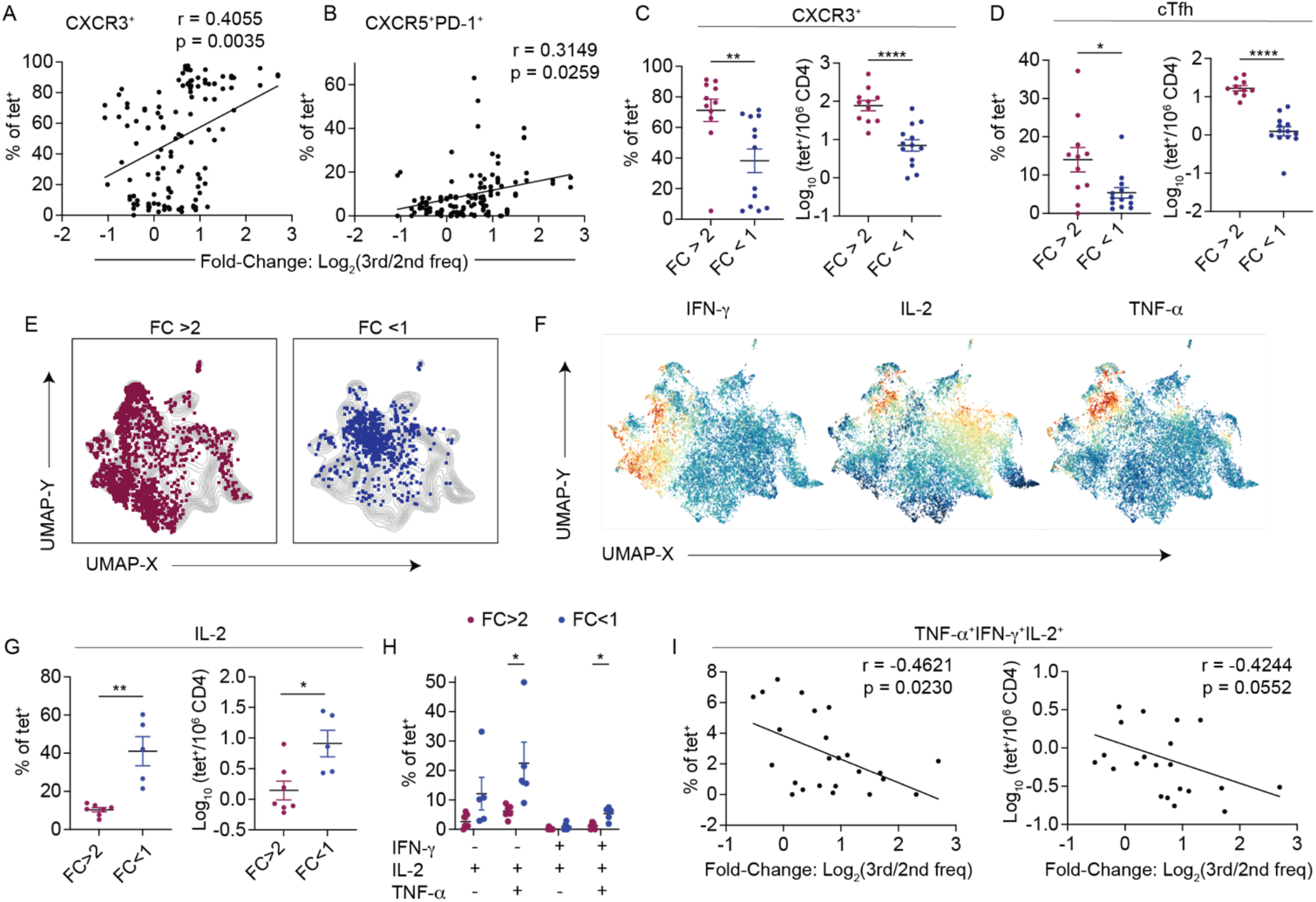
Response to booster is associated with distinct functional states. (A–B) Correlation between fold change from post-2^nd^ dose baseline to post-3^rd^ dose and the proportion of CXCR3^+^ (A) or cTfh subset (B) within tetramer^+^ populations. Spearman correlation was performed. (C–D) Comparison of CXCR3^+^ (C) and cTfh (D) proportions and frequencies between populations with fold change >2 or <1 from post-dose 2 to 3. Percentages: Mann–Whitney test; Frequencies: Welch’s t-test. (E) PBMC were stimulated with PMA and ionomycin for 6 hours, followed by tetramer staining and enrichment using post-3^rd^ dose blood from 7 donors, followed by surface and intracellular antibody staining and analyses on spectral cytometry. Populations were stratified based on booster-dependent fold change and grouped as FC > 2 (n = 7 populations, 2141 cells) or FC < 1 (n = 5 populations, 849 cells). UMAPs show tetramer^+^ cells from each group projected onto all post–third dose SARS-CoV-2–specific CD4^+^ T cells (n = 24 populations, 15,432 cells). (F) UMAPs display the staining intensity of the indicated markers. Data include 15,432 cells from 24 tetramer^+^ populations. (G) IL-2 percentage and frequency within tetramer^+^ populations in the FC > 2 (n = 7) versus FC < 1 groups (n = 5) after the 3^rd^ vaccine dose. Mann-Whitney test was used. (H) Cytokine combinations in IL-2^+^ subset, shown as percentage of tetramer^+^ cells by FC > 2 or < 1. Multiple Mann-Whitney tests were performed. (I) Correlation between booster-induced fold change and the percentages (left) or frequencies (right) of TNF-α^+^IFN-γ^+^IL-2^+^ tetramer^+^ cells (n = 24). Percentages: Spearman correlation; Frequencies: Pearson correlation. Each symbol represents the mean for a tetramer^+^ population, averaged across 1-5 independent experiments. Data are represented as mean ± SEM. *p < 0.05, ** p< 0.01, and ****p < 0.0001.

### Vaccination by the ancestral spike vaccination expands CD4^+^ T cell memory to spike variants

As a rapidly evolving pathogen, SARS-CoV-2 poses a continual risk of immune escape. Although many SARS-CoV-2 epitopes are conserved across variants, it remains unclear how effectively CD4^+^ T cells recognize mutated sequences. To investigate this question, we generated tetramers for three peptide pairs consisting of a vaccine-matched sequence and the same sequence carrying a point mutation found in a viral variant (Fig. 7A). Staining with Wu-1 and variant peptide-loaded tetramers was performed on blood collected before and after vaccination to distinguish ancestral-specific, variant-specific, and cross-reactive CD4^+^ T cells that bound both tetramers (Fig. 7B). As observed for Wu-1 epitopes, vaccination elicited robust CD4^+^ T cell responses to variant spike sequences, including a substantial population that recognized both mutated and unmutated spike sequences (Fig. 7B-C). To define the composition of this response, Wu-1 and the corresponding variant tetramer^+^ cells were stratified into Wu-1-only, variant-only, and double-positive subsets. At the pre-vaccination baseline, each subset was present at similar frequencies (Fig. 7D). Following vaccination, however, cross-reactive cells preferentially expanded and become the dominant subset in percentage and frequency compared to singly binding tetramer^+^ cells (Fig. 7E-F). Consistent with affinity-based selection, cross-reactive T cells detected after the 3^rd^ vaccine dose showed a greater increase in tetramer MFI from baseline compared with cells binding only Wu-1 or variant tetramers (Fig. 7G-H). To evaluate functional response, individual S690/S690_S704L double tetramer^+^ CD4^+^ T cells were single-cell sorted from HD11’s post–3^rd^ vaccine time point into 96-well plates and expanded in vitro. A total of 31 clones were generated, all of which responded to stimulation with S690 peptide (Fig. 7I, S6A-B). Six of the most robustly proliferating clones were selected for further analyses and stimulated with decreasing concentrations of S690 and S690_S704L peptides. All six clones produced TNF-α in response to both peptides, but responses to the variant sequence were generally weaker, as indicated by higher average EC50 values compared with the ancestral sequence (Fig. 7J, S6C). Collectively, these data demonstrate that COVID-19 mRNA vaccines and boosters containing ancestral spike elicit cross-reactive T cells that can recognize and respond to viral variants.

**Figure 7.**
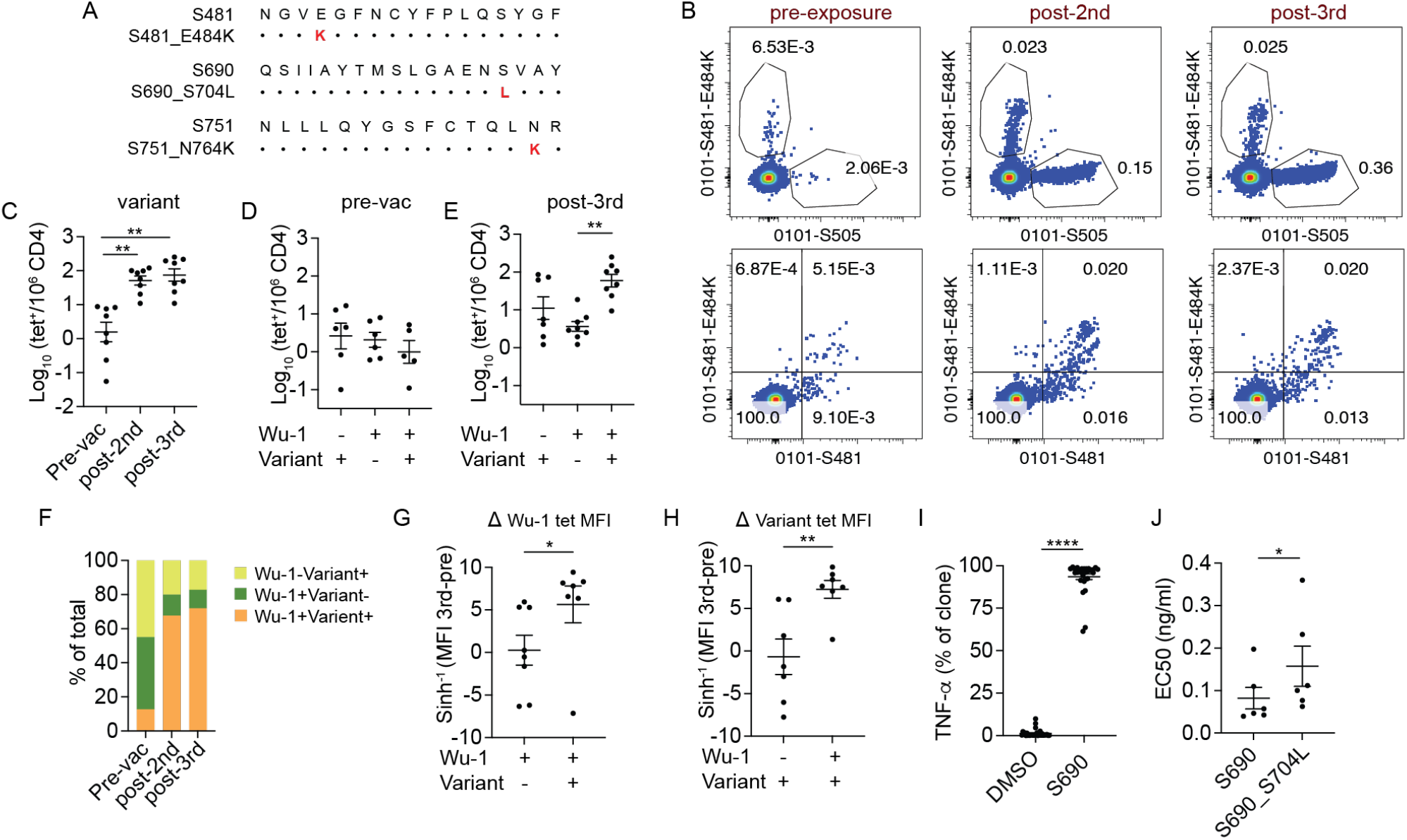
Ancestral spike vaccination expands cross-reactive T cells recognizing viral mutations. (A) Sequence alignment of three Wu-1 and variant peptide pairs. Red highlights amino acid differences. (B) Representative direct ex vivo tetramer staining of Wu-1 and variant tetramer^+^ CD4^+^ T cells before vaccination, post-2^nd^, and post-3^rd^ vaccine doses. C) Frequency of variant tetramer^+^ CD4^+^ T cells across time points (n = 8). RM one-way ANOVA with Tukey multiple comparison test was performed. (D-E) Pre-vaccination (D) and post-3^rd^ dose (E) frequencies of single-reactive (Wu-1 or variant) and cross-reactive (Wu-1^+^Variant^+^) tetramer^+^ populations. Kruskal-Wallis test with Dunn’s multiple comparison test were performed. (F) Relative abundance of single-reactive (Wu-1 or variant-single), and cross-reactive (Wu-1^+^Variant^+^) tetramer^+^ populations over time. (G-H) Changes in tetramer MFI after the 3^rd^ vaccine dose relative to pre-vaccine baseline for cross-reactive versus single Wu-1 (G) or variant (H) tetramer^+^ populations. Mann-Whitney tests were used. (I) Plot summarizes TNF-α production by 31 single-cell derived S690 T cell clones after 5-hour stimulation with monocyte-derived dendritic cells treated with DMSO vehicle control or S690 peptide. Wilcoxon test was performed. (J) T cell clones were stimulated with decreasing concentrations of S690 or S690_S704L peptides. The response was measured by TNF-α production and quantified by EC50 values. Data are represented as mean ± SEM. *p < 0.05, ** p< 0.01, and ****p < 0.0001.

## Discussion

We longitudinally tracked human spike-specific CD4^+^ T cell responses to COVID-19 mRNA vaccination, from the pre-vaccine state through post-booster, to define how primary memory is established and remodeled by antigen exposures. Ex vivo pMHC tetramer staining and enrichment enabled direct quantification of population size and differentiation states. We captured both dominant and subdominant CD4^+^ T cell specificities, providing a comprehensive view of vaccine-induced memory dynamics. Analyses of the precursor repertoire showed a heterogeneous baseline that reflected imprints of prior exposures ^20,30,31^. Although antigenically related pathogens could theoretically provide additional sources of cross-reactive antigens that prime and deplete the naïve repertoire, our data does not support this idea. Comparison of Wu-1 spike- and YFV-specific T cells showed similar precursor and naïve subset frequencies, indicating that the naïve repertoire remains largely intact despite antigenically related exposures. Instead, YFV and SARS-CoV-2 vaccine responses differed in their relationship to pre-existing memory. For YFV, populations with large pre-existing memory were numerically disadvantaged after vaccination, whereas spike-specific responses lacked the clear inverse relationship with baseline memory frequency. Higher pre-vaccine memory frequencies were associated with a greater abundance of the same tetramer^+^ population after COVID-19 vaccination, suggesting that vaccines can more effectively leverage pre-formed memory cells generated by closely related antigens.

After the initial vaccinations, we observed a substantial increase in antigen-specific memory and a reorganization of the immunodominance hierarchy. Vaccination selectively recruited a small subset of high avidity precursors, which becomes the dominant populations in the memory repertoire. These findings are consistent with past studies on YFV and COVID-19 vaccines ^19,32^. The ability of COVID-19 vaccination to drive appropriate T cell selection likely contributes to its protective efficacy, as failure to generate high-avidity CD8^+^ T cells has been associated with HIV vaccine failure and with severe SAR-CoV-2 infection ^28,33^. After the initial vaccinations, many individuals received booster doses, reflecting a broader pattern of repeated antigen exposures in human populations. A major unresolved question is how a booster vaccine influences T cell responses. Unlike antibodies, which continue to rise after boosting, T cell responses appear to plateau after the second mRNA vaccine dose in peptide stimulation assays ^11^, calling into question of the benefit of the booster. Our epitope-level analyses uncovered a dynamic landscape of booster responses. While the immunodominance was largely set after the second vaccine dose, with no significant change in tetramer MFI to suggest major additional affinity-based selection, the 3^rd^ vaccine dose refined the dominance hierarchy. Populations that responded to boosting were maintained or increased in rank, whereas non-expanding populations declined. In addition, the 3^rd^ vaccine dose fine tunes the quality and fate of CD4^+^ T cell memory, which are critical for effective protective immunity. Following the first mRNA COVID-19 vaccine, early Th1 and Tfh responses correlated with subsequent CD8^+^ T cell and neutralizing antibody responses ^34^. Here, we showed that these key CD4^+^ T cell differentiation states diverge with the magnitude of the primary memory response and are reinforced by boosting. Dominant, highly expanded memory adopted a Th1-biased program, whereas less-expanded memory preferentially acquired Tfh differentiation. Boosting selectively promoted further Th1 and Tfh differentiation among subdominant populations. The effects of the booster were heterogeneous, with some failing to expand. These were generally less abundant populations that retained high polyfunctional potential. While vaccines often aim to elicit immunodominant populations, subdominant responses can acquire effector functions and provide protective immunity ^35-38^. In some models, they outperform dominant responses and control viral replication without driving immunopathology ^37^. These ‘booster resistant’ populations may represent a potential reservoir of untapped immunity, and targeting them in future vaccine design could broaden the functional diversity of T cell response and enhance vaccine efficacy.

In addition to memory T cells targeting the vaccine-matched spike, our data demonstrated that COVID-19 mRNA vaccines encoding the ancestral spike also generated a pool of high-avidity, cross-reactive T cells capable of recognizing future variants. The emergence of variants that diverge from the original vaccine sequence poses a major challenge for many vaccines and was a particular concern for COVID-19 during the pandemic. Unlike neutralizing antibodies, which are less efficacious against mutated viruses, T cells can target conserved internal viral proteins and this helps to maintain their responses across viral variants ^7-10^. Our findings extend key insights from these prior studies by resolving responses at the epitope level. They indicate that the maintenance of T cell responses to SARS-CoV-2 variants reflects not only substantial sequence conservation but also an intrinsic capacity of TCRs to tolerate certain mutations. Moreover, cross-reactive T cells showed a selective advantage and were preferentially recruited over mono-reactive subsets. We speculate that broadening of T cell responses may buffer against pathogen escape but could also increase the risk of off-target recognition and autoimmunity in susceptible individuals.

In conclusion, our data highlight the dynamic nature of T cell memory and the layered impact of repeat exposures on CD4^+^ T cell responses. By linking pre-existing repertoires to memory established after primary vaccination and boosting, our data reveal population-specific dynamics and divergent differentiation programs. These epitope-level insights advance understanding of how the human memory repertoire is established, refined, and highlight opportunities for future vaccine strategies.

## Limitations of study

This study has several limitations. The cohort was small and limited to young-to middle-aged adults. Larger studies across diverse ages and baseline immune statuses are needed to capture human heterogeneity in vaccine-induced memory responses. Although our experiments reveal distinct T cell fates and functional potentials among vaccine-responding populations, considerable phenotypic and functional heterogeneity likely exists beyond the subsets analyzed. Early expansion and contraction dynamics were not examined. Our study is confined to blood. Tissue-resident and lymphoid populations may exhibit different types of responses. Finally, a limited number of ancestral-variant peptide pairs was examined. Further studies are needed to define the full breadth and cross-reactive potential of COVID-19 vaccine-induced CD4^+^ T cell memory.

## Acknowledgments

We thank our study subjects for their participation and members of Hensley lab for technical assistance with serological assays.

## Author contributions

Conceptualization, L.F.S.; Experimentation, Y. L., X.S., A.A, H.C., E.M.D; Computational analyses, Y.L., A.O.S, J.J.; Study recruitment: S.A., A.S., H.J.; Modeling and statistical support, W.B.; Supervision, S.H., L.F.S.; Manuscript preparation, L.F.S., Y.L.

## Competing interests

None

### VA Disclaimer

The views expressed in this article are those of the authors and do not necessarily reflect the position or policy of the Department of Veterans Affairs or the United States government.

During the preparation of this work, L.F.S. used ChatGPT for grammar correction and to improve readability. After using this tool, the author reviewed and edited the content as needed and take full responsibility for the content of the published article.

## Supplementary Material

**Figure S1:**
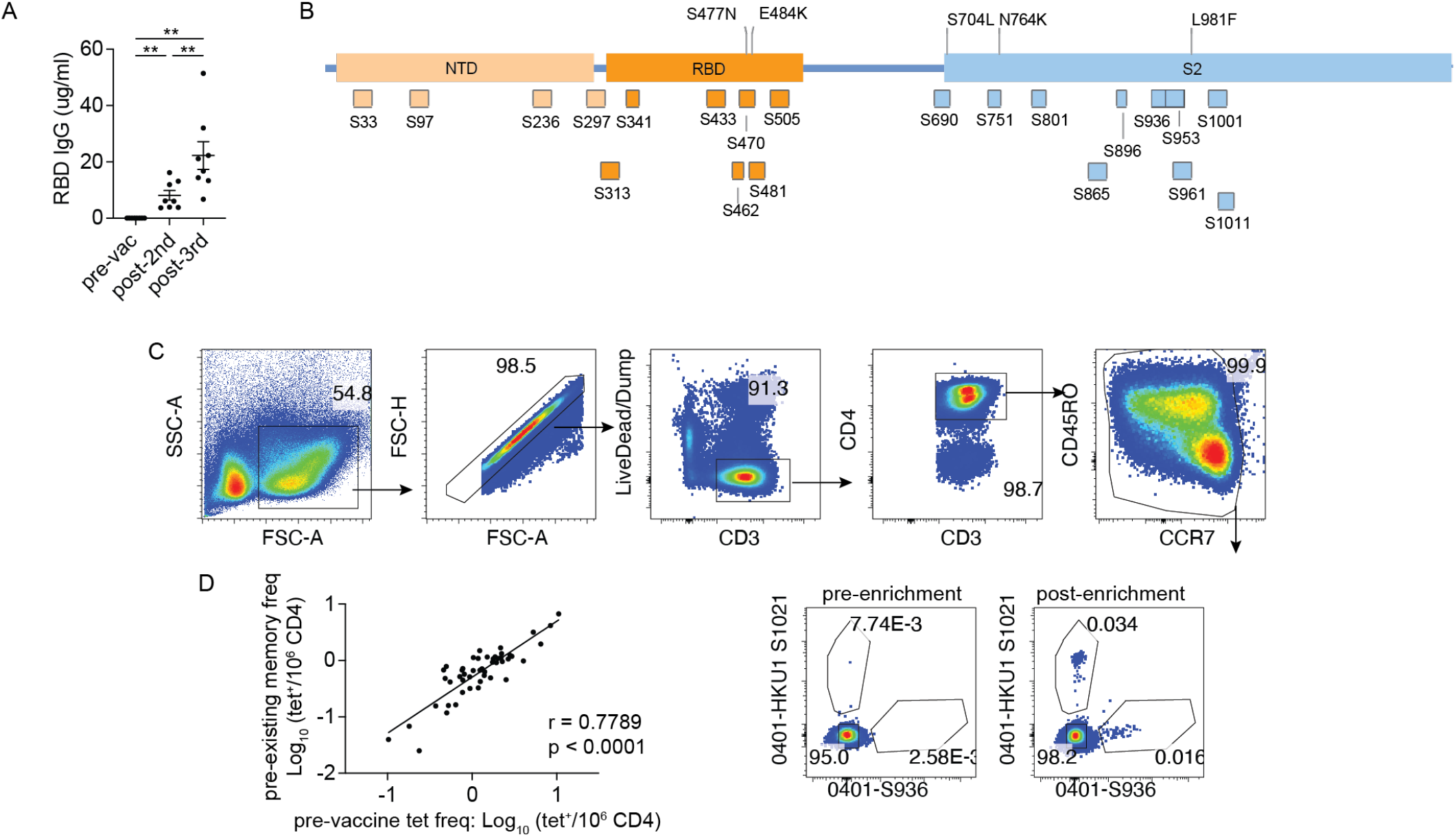
SARS-CoV-2 epitopes and gating strategies. (A) RBD-specific IgG levels measured in plasma collected at the indicated time points. RM one-way ANOVA with Holm-Sidak multiple comparison test were performed. (B) Schematic of spike protein showing the position of peptides used in direct ex vivo tetramer analysis. (C) Representative plots show the gating strategy for identifying tetramer^+^ cells. (D) Correlation between the frequency of tetramer^+^ T cell and pre-existing memory T cells within each population in the pre-vaccine samples (n = 50). Association was measured by Spearman correlation.

**Figure S2:**
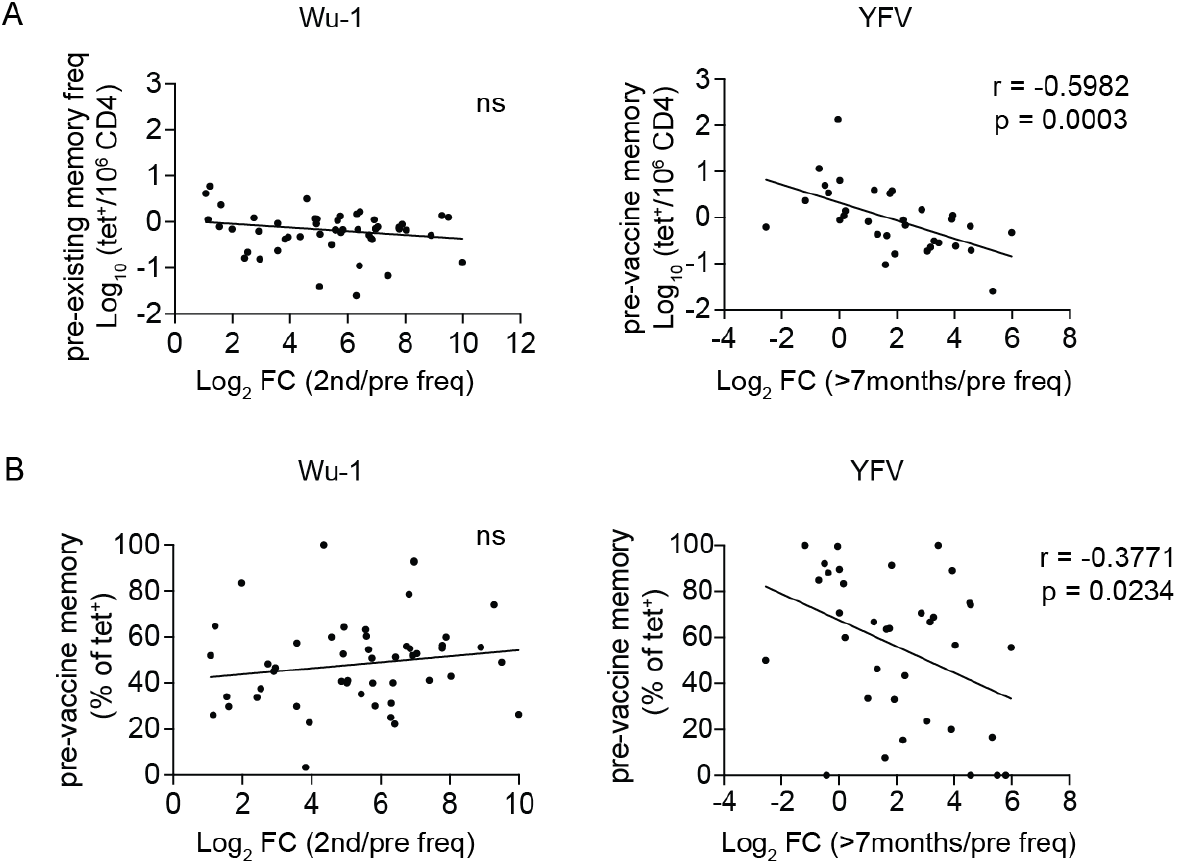
Baseline Memory Differentially correlate with YFV and COVID-19 Vaccine Responses. (A-B) Relationship between the gain in tetramer^+^ T cells after the two-dose COVID-19 vaccines (left) or YFV vaccine (right) and pre-existing memory, as frequency (A) or a percentage within tetramer^+^ cells (B) in the pre-vaccine samples. Pearson or Spearman correlation was used depending on Shapiro-Wilk test for normality.

**Figure S3:**
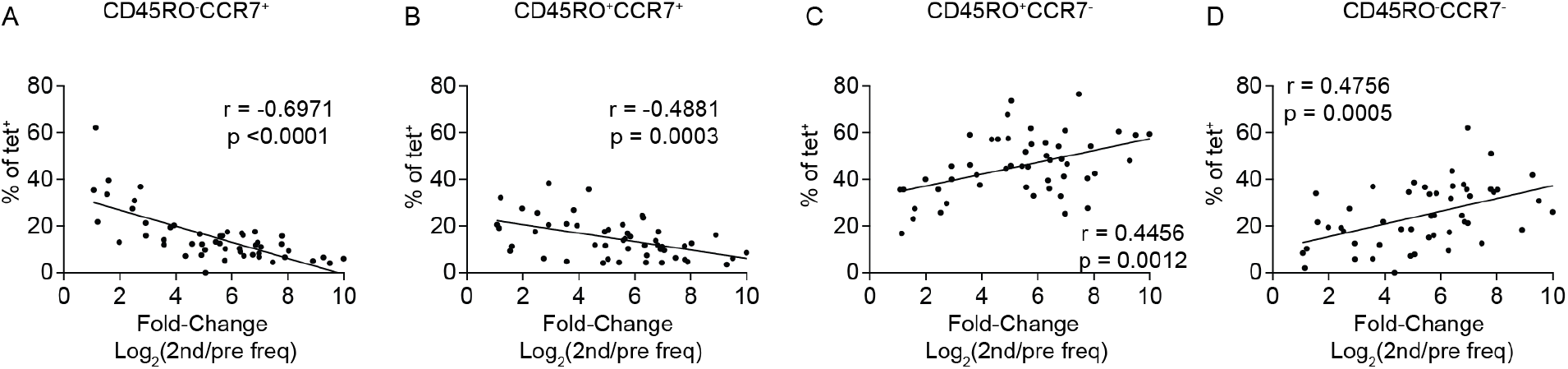
Fold expansion after vaccination is associated with distinct memory phenotypes. (A–D) The percentage of naïve-like (A), CM (B), EM (C), and TEMRA (D) subsets within tetramer^+^ populations correlates with vaccine-induced fold change after the 2^nd^ dose (n = 50). Pearson or Spearman correlation was used depending on Shapiro-Wilk test for normality.

**Figure S4:**
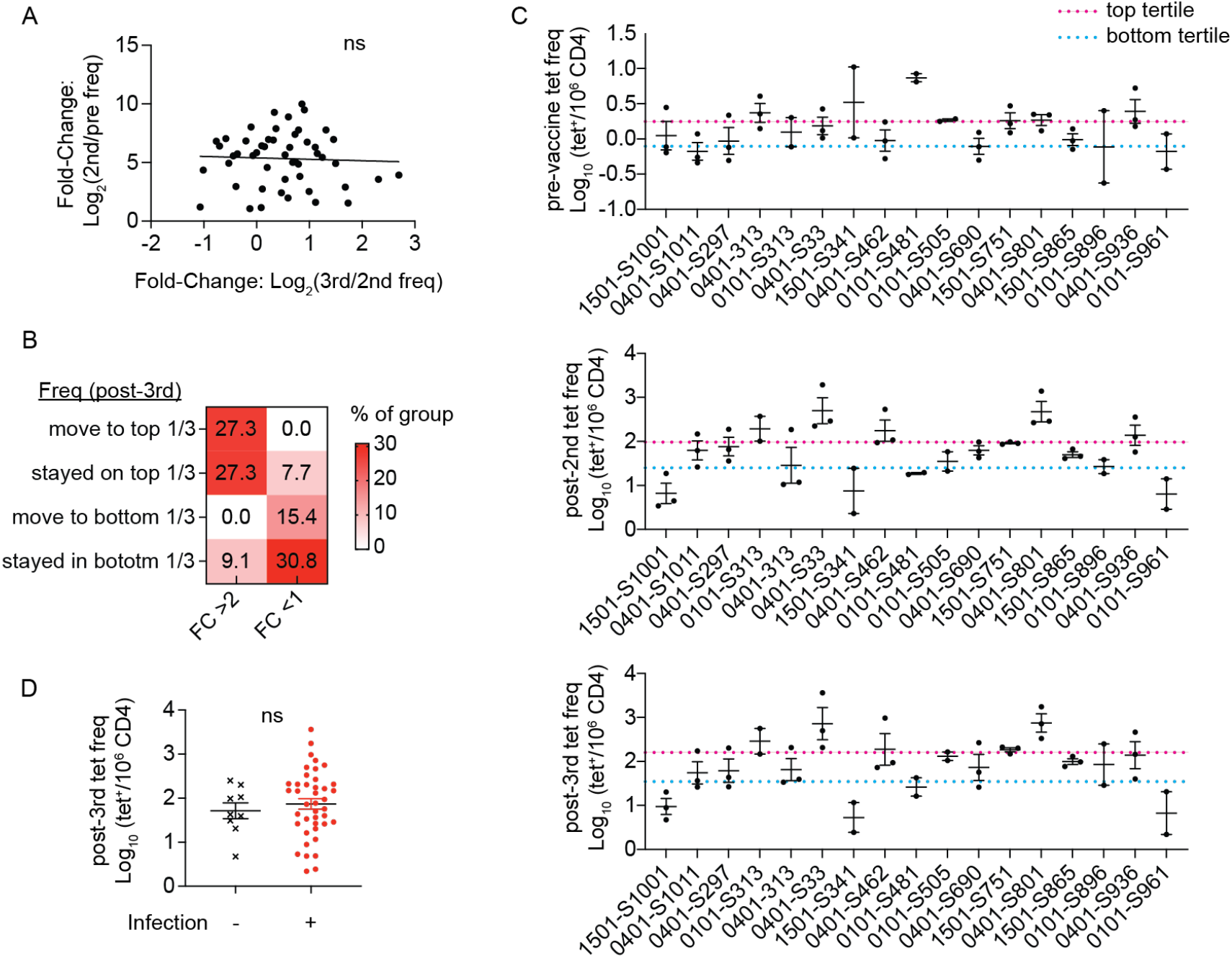
The magnitude of vaccine responses by specificity and in relationship to prior exposures. (A) Correlation between responses to the initial 2-dose series (post-2^nd^/pre) and responses to the booster (post-3^rd^/post-2^nd^) (n = 50). Pearson correlation was performed. (B) Percentage of FC > 2 or FC < 1 tetramer^+^ populations that stayed or moved into the top or bottom one-third of frequencies after the third vaccine dose (FC > 2 n= 11, FC <1 n = 13). (C) Plot shows tetramer^+^ populations for epitopes quantified in at least two individuals. Each dot represents a different donor for that specificity. Populations were ranked by frequency and divided into tertiles. Differences in the distribution of populations across tertiles of responses were assessed using the Fisher’s Exact test: not significant pre-vaccination, p < 0.001 post-2^nd^ dose, p = 0.009 post-3^rd^ dose. (D) Frequencies of Wu-1 tetramer^+^ populations after the booster, stratified by donor infection status between the 2^nd^ and 3^rd^ vaccine doses (uninfected, n = 9; infected, n = 41). Welch’s t-test as performed.

**Figure S5:**
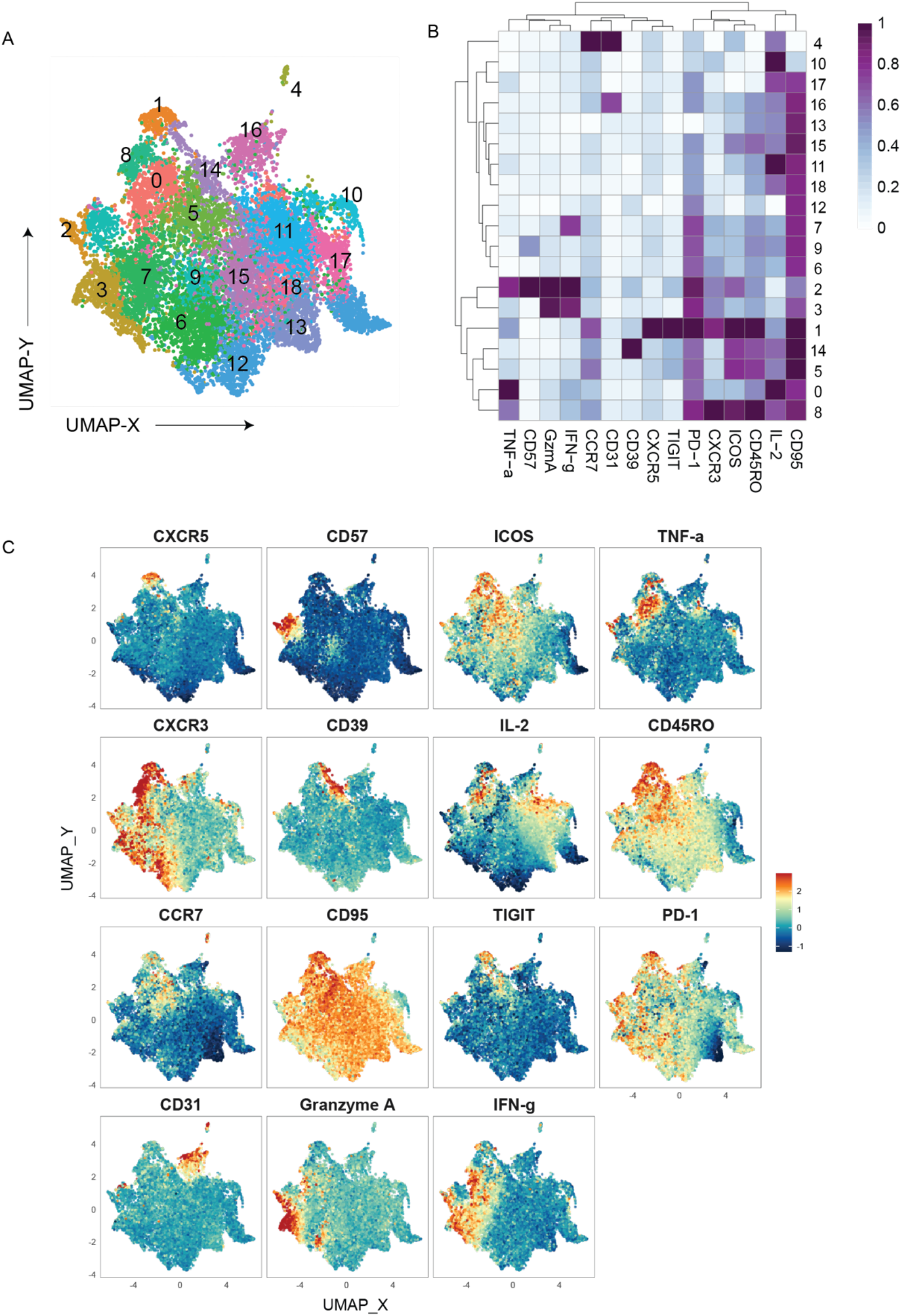
Phenotypic landscape of SARS-CoV-2–specific CD4^+^ T cells after booster. (A) UMAP displays phenograph-defined clusters of Wu-1–specific CD4^+^ T cells. (B) Heatmap shows the median staining signal of individual markers for clusters shown in (A). (C) UMAPs display the staining intensity of the indicated markers. Data include 15,432 cells from 24 tetramer^+^ populations in post–3rd dose blood samples from 7 individuals.

**Figure S6:**
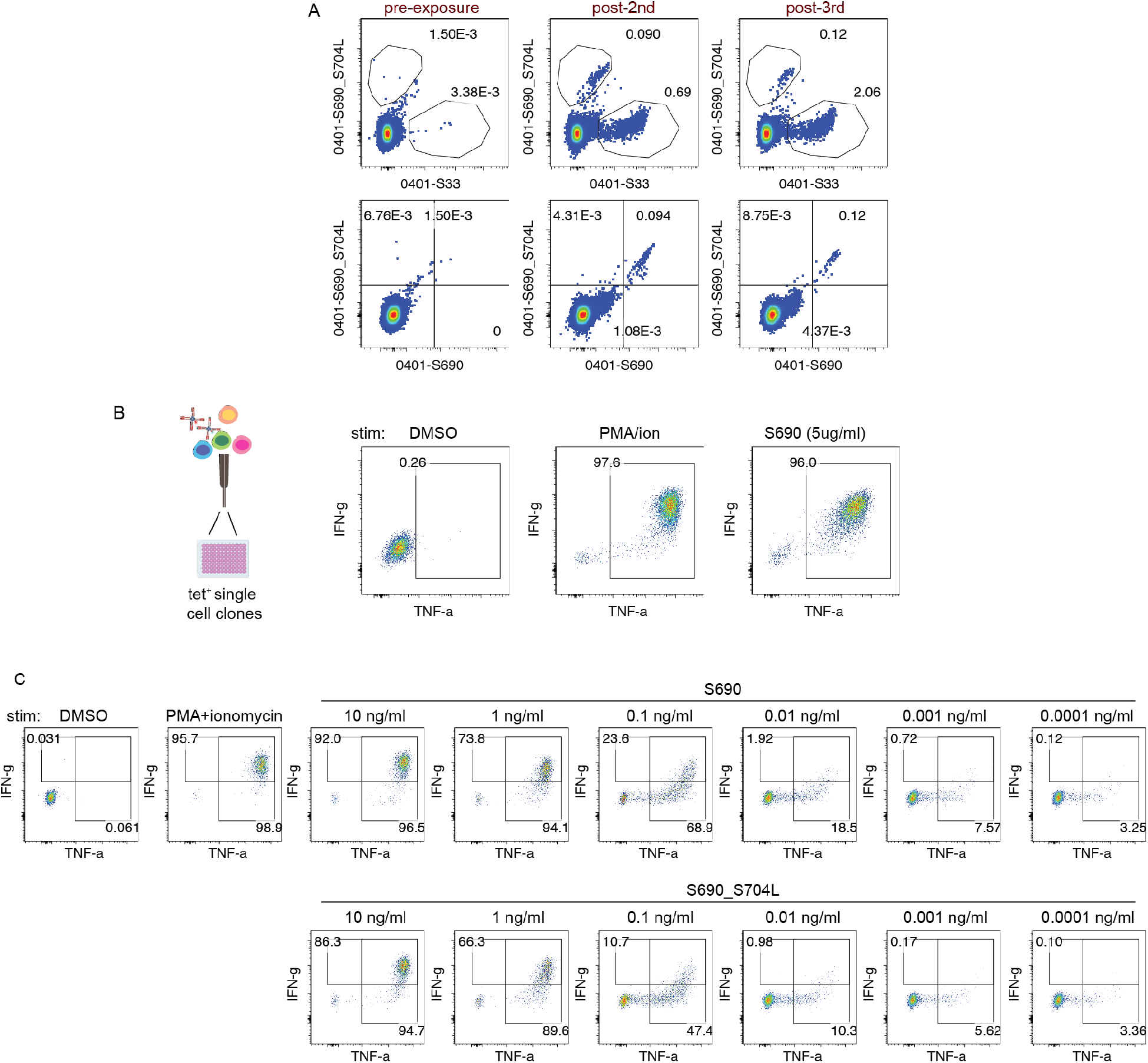
Cross-recognition of variant sequences by Wu-1 tetramer^+^ CD4^+^ T cells. (A) Representative direct ex vivo tetramer staining of S690, S690_S704L, and unrelated spike–specific CD4^+^ T cells before vaccination, after the 2nd dose, and after the 3rd COVID-19 vaccine dose. (B) Plots show cytokine production by a representative single-cell derived S690 T cell clones after 5-hour stimulation with monocyte-derived DCs treated with DMSO vehicle control, PMA and ionomycin, or S690 peptides. (C) Representative cytokine responses to DCs loaded with decreasing concentrations of S690 or S690_S704L peptides.

**Table S1:** Donor information.

| ID | Age | Sex | Vaccine: 1st and 2nd dose |  | Vaccine: 3rd dose |  | COVID-19 infection | pre-vaccine blood | post-vaccine blood |  |
| --- | --- | --- | --- | --- | --- | --- | --- | --- | --- | --- |
|  |  |  | Name | Date | Name | Date |  | pre-1st dose (days) | post-2nd (days) | post-3rd (days) |
| D01 | 54 | Female | Moderna mRNA-1273 | 3/1/21, 3/25/21 | Moderna mRNA-1273 | 11/8/21 | 5/24/22 | -123 | 186 | 241, 332 |
| D03 | 22 | Female | Moderna mRNA-1273 | 2/25/21, 3/25/21 | Moderna mRNA-1273 | 12/7/21 | 3/14/22 | -2 | 256 | 197 |
| D04 | 24 | Female | Pfizer BNT162b2 | 3/22/21, 4/9/21 | Moderna mRNA-1273 | 1/5/22 | 12/27/22 | -32 | 213 | 197 |
| D05 | 32 | Male | Pfizer BNT162b2 | 3/19/21, 4/9/21 | Pfizer BNT162b2 | 11/30/21 | 4/11/22 | -31 | 199 | 198 |
| D09 | 36 | Female | Pfizer BNT162b2 | 3/15/21, 4/6/21 | Pfizer BNT162b2 | 2/3/22 | NA | -32 | 252 | 204 |
| D10 | 32 | Female | Pfizer BNT162b2 | 4/9/21, 4/30/21 | Pfizer BNT162b2 | 11/3/21 | 12/28/21 | -23 | 182 | 187 |
| D11 | 34 | Male | Moderna mRNA-1273 | 4/13/21, 5/11/21 | Moderna mRNA-1273 | 1/10/22 | 12/22/21 | -19 | 205 | 172 |
| D21 | 29 | Female | Pfizer BNT162b2 | 5/1/21, 5/24/21 | Pfizer BNT162b2 | 1/3/22 | NA | -3 | 176 | 190 |

**Table S2:**
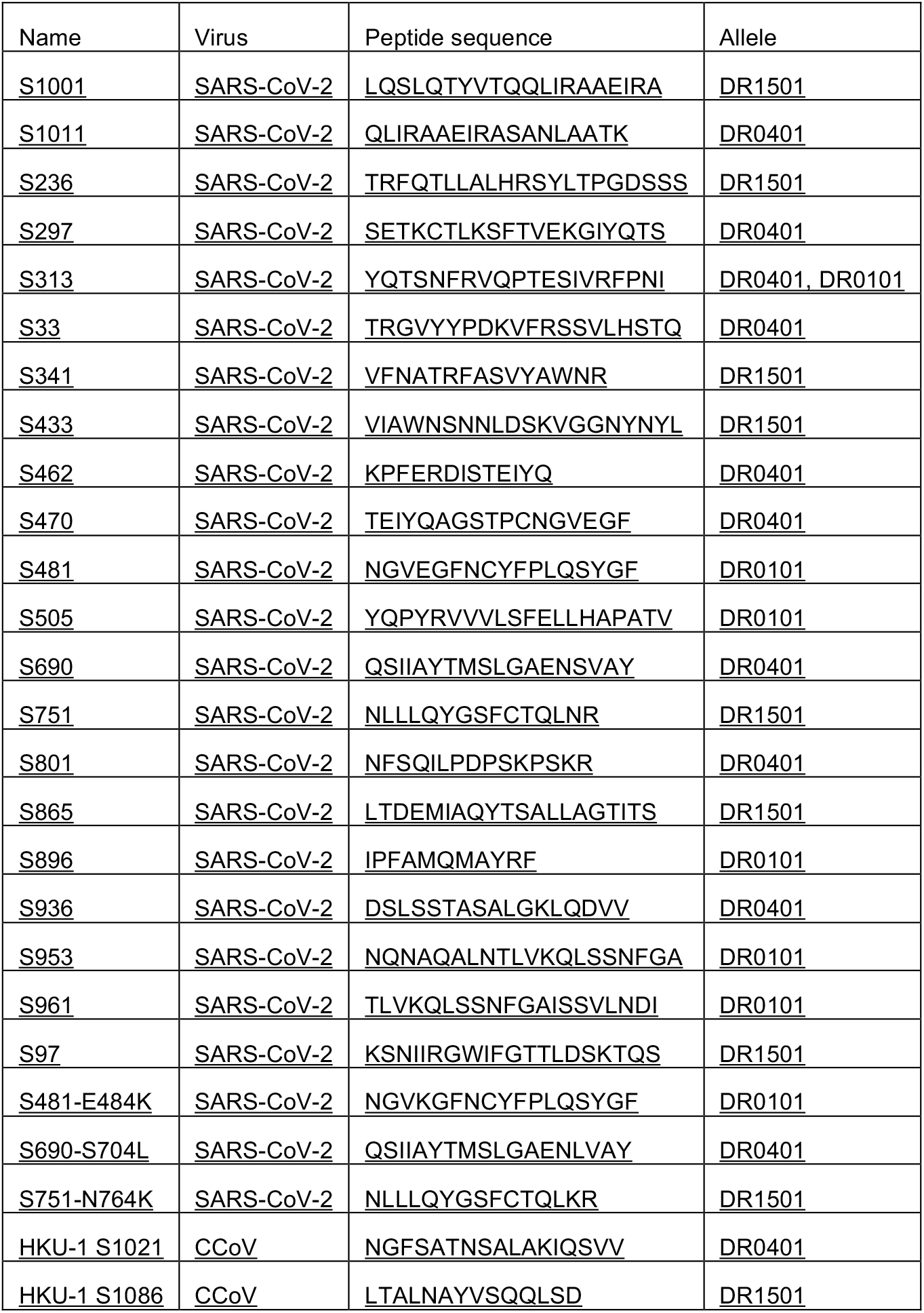
List of peptides.

**Table S3:** List of antibodies.

| Antibodies | Vendor | Identifier |
| --- | --- | --- |
| Mouse anti-human CD45RO antibody | BioLegend | Cat# 304238; RRID: AB_2562153 or Cat#304226; RRID: AB_2563818 |
| Mouse anti-human CD197 (CCR7) antibody | Biolegend | Cat# 353234; RRID: AB_2563867 |
| Mouse anti-human CD4 antibody | Biolegend | Cat# 344634; RRID: AB_2566017 or Cat# 317448; RRID:AB_2565847 or Cat# 317410; RRID:AB_571955 |
| Mouse anti-human CD4 antibody | BD Biosciences | Cat# 612962; RRID: AB_2739452 |
| Mouse anti-human CD3 antibody | Biolegend | Cat# 300410; RRID: AB_314064 or Cat# 317322; RRID:AB_2561911 |
| Mouse anti-human CD3 antibody | BD Biosciences | Cat# 612895; RRID:AB_2870183 |
| Mouse anti-human CD278 (ICOS) antibody | BD Biosciences | Cat# 750321; RRID: AB_2874511 |
| Mouse anti-human CD11b antibody | Biolegend | Cat# 301342; RRID: AB_2563395 |
| Mouse anti-human CD19 antibody | Biolegend | Cat# 363010; RRID: AB_2564193 |
| Mouse anti-human TNF-α antibody | eBioscience | Cat# 367-7349-42; RRID: AB_2896048 or Cat# 502930; RRID:AB_2204079 |
| Rat anti-human IL-2 antibody | Biolegend | Cat# 500338; RRID: AB_2562632 |
| Mouse anti-human CD279 (PD-1) antibody | Biolegend | Cat# 329968; RRID: AB_2894474 or Cat# 329920; RRID: AB_10960742 |
| Rat anti-human CD185 (CXCR5) antibody | BD Biosciences | Cat# 564624; RRID: AB_2738871 |
| Mouse anti-human CD183 (CXCR3) antibody | Biolegend | Cat# 353738; RRID: AB_2565924 or Cat# 353716; RRID: AB_2561448 |
| Mouse anti-human CD8a antibody | BD Biosciences | Cat#366-0088-42; RRID: AB_2920986 |
| Mouse anti-human Granzyme A antibody | Biolegend | Cat# 507207; RRID: AB_439755 |
| Mouse anti-human IFN-γ antibody | Biolegend | Cat# 502552; RRID: AB_2888716 or Cat# 502506; RRID:AB_315231 |
| Mouse anti-human CD57 antibody | BD Biosciences | Cat#567621; RRID: AB_3683865 |
| Mouse anti-human CD39 antibody | BD Biosciences | Cat#746454; RRID: AB_2743756 |
| Mouse anti-human CD95 antibody | Biolegend | Cat#305644; RRID: AB_2632623 |
| Mouse anti-human TIGIT(VSTM3) antibody | Biolegend | Cat#372736; RRID: AB_2892455 |
| Mouse anti-human CD31 antibody | Biolegend | Cat#303154; RRID: AB_3097292 |

## Lead Contact

Further information and requests for resources and reagents should be directed to and will be fulfilled by the Lead Contact, Laura F. Su

## Material Availability

This study did not generate new unique reagents.

## Data and Code Availability

Data were analyzed uses existing computational packages.

## EXPERIMENTAL MODEL AND SUBJECT DETAILS

### Study design

Volunteers were screened for prior SARS-CoV-2 exposure by questionnaire and serologic testing. All donors underwent high-resolution HLA typing (Histogenetics, NY). Eight participants with no known prior SARS-CoV-2 exposure, negative spike RBD serology at baseline, and carry HLA-DRB1*010, 0401, or 1501 alleles were enrolled. Blood was collected by Leukapheresis prior to vaccination. Post-vaccine blood samples were obtained by leukapheresis or peripheral blood draws approximately 6 - 8 months after completion of the primary mRNA vaccine series, and approximately 6 – 11 months following the third mRNA dose. All participants received mRNA-based SARS-CoV-2 vaccines for both the primary series and booster immunizations (BNT162b2 or mRNA-1273). All samples were de-identified and obtained with IRB regulatory approval from the University of Pennsylvania (protocol 820884). Subject characteristics are shown in Table S1.

### Sex as a biological variable

Sex was considered by including both males and female study participants.

### Primary cells

#### Human samples preparation

Plasma was isolated from whole blood by centrifugation and stored at −80°C. PBMC was isolated by Ficoll-Paque (GE Healthcare) density gradient centrifugation according to the manufacturer’s protocol. T cells and monocytes were enriched from leukapheresis products by negative selection using RosetteSep (STEMCELL Technologies). Isolated cells were cryopreserved in fetal bovine serum supplemented with 10% DMSO and stored in liquid nitrogen until use.

#### Cell lines

Hi5 cells (ThermoFisher) were maintained by insect cell culture medium (Lonza Bioscience) supplemented with 0.02% gentamicin at 28°C.

### SARS-CoV-2 ELISA

Antibodies specific to the receptor binding domain (RBD) of the SARS-CoV-2 spike protein were measured by enzyme-linked immunosorbent assay (ELISA) as described previously ^39^. The plasmid encoding RBD was gifted by Florian Krammer (Mt. Sinai) and purified using nickel-nitrilotriacetic acid resin (Qiagen). ELISA plates (Immulon 4 HBX, Thermo Fisher Scientific) were coated with 2mg/mL recombinant RBD or PBS and incubated overnight at 4ºC. The plates were then washed with phosphate-buffered saline containing 0.1% Tween-20 (PBS-T) before blocking with PBS-T supplemented with 3% non-fat milk powder for one hour. Heat-inactivated plasma samples (56ºC for one hour) were diluted in buffer containing PBS-T and 1% non-fat milk powder. The ELISA plates were then washed with PBS-T, diluted samples were added to each well, and the plates were incubated for two hours. The plates were washed with PBS-T and incubated for 1 hour with 1:5000 diluted goat anti-human IgG-HRP (Jackson ImmunoResearch Laboratories). The plates were washed once more with PBS-T before adding SureBlue 3,3’,5,5’-tetramethylbenzidine substrate (KPL) and incubating for five minutes. 250mM hydrochloric acid was added to stop the reaction and the plates were read on a SpectraMax 190 microplate reader (Molecular Devices) at an optical density (OD) of 450 nm. Each plate also included the monoclonal antibody CR3022, which was used to convert OD values into relative antibody concentrations. The plasmids used to express CR3022 were provided by Ian Wilson (Scripps).

### Epitope identification and tetramer production

Spike protein sequences of SARS-CoV-2 and seasonal human common cold coronavirus (CCoVs) were obtained from the National Center for Biotechnology Information (NCBI) database (SARS-CoV-2 spike, YP_009724390.1; HKU1 spike, ABD96198.1). Spike sequences for SARS-CoV-2 variants (Alpha, Beta, Gamma, Delta, Mu, BA.1, BA.2, BA.2.12.1, BA.4/BA.5, XBB1.5, and KP.2) were obtained from the outbreak.info database ^40^. Peptides candidates included sequences identified in previous studies and predicted to bind HLA-DRB1*0101, 0401 and1501 with a consensus percentile under 20 using NetMHCIIpan 4.1 from IEDB analyses resource ^17,18^. PBMC obtained after vaccination were stimulated with candidate peptides at 0.4ug/ml per peptide for 3 weeks and stained with tetramers to evaluate T cell expansion. Peptides that elicited robust responses were validated by direct ex vivo tetramer staining (Table S2). For tetramer production, HIS-tagged HLA-DRA/B1*0101, 0401, and 1501 protein monomers were produced by Hi5 insect cells and purified from culture supernatant using Ni-NTA (Qiagen), followed by size exclusion chromatography (AKTA, Cytiva). Protein biotinylation, peptide exchange, and tetramerization were performed using standard protocols as previously described ^19,20,41,42^. In brief, HLA-DR proteins were incubated with thrombin (Millipore) at room temperature for 3 - 4 hours and exchanged with peptides of interest in 50-fold excess at 37°C for 16 hours. Peptide-loaded HLA-DR monomers were incubated with fluorochrome-conjugated streptavidin at 5:1 ratio for 2 min at room temperature, followed by a 15 min incubation with an equal volume of biotin-agarose slurry (Millipore). Tetramers were exchanged into PBS, concentrated using Amicon ULTRA 0.5ml 100KDa (Millipore), and kept at 4 °C for no more than one week prior to use.

### Direct ex vivo T cell analyses and cell sorting

Pre-vaccination analyses were performed using 30–100 million CD3 or CD4 enriched T cells. For post-vaccination analyses, 10–30 million cells were used. Tetramer staining and enrichment was performed as previously described, with up to four tetramers per reaction ^19,20,41,42^. Tetramer-positive cells were enriched using anti-fluorochrome and anti-HIS MicroBeads followed by enrichment through LS columns according to manufacturer protocol (MiltenyiBiotec). For antibody staining, tetramer enriched samples were stained with viability dye, exclusion markers (anti-CD19 and anti-CD11b), and surface antibodies (anti-CD3, anti-CD4, anti-CD45RO, anti-CCR7, anti-CXCR5, anti-CXCR3, and anti-PD-1, Tabls S3) for 45 min at 4°C. Cells were then fixed in 2% paraformaldehyde, acquired on an LSR II flow cytometer (BD), and analyzed using FlowJo software (BD). Frequencies were calculated by mixing one-tenth of each sample with 200,000 fluorescent counting beads (Spherotech) for normalization ^42^. For longitudinal analyses, samples collected at pre- and post-vaccine time points were performed in the same experiment to minimize technical variability. For single-cell cloning, tetramer-enriched cells were prepared as above without fixation and sorted using a FACSAria II (BD).

### T Cell Stimulation and Intracellular Cytokine Staining

Cryopreserved T cells were thawed, rested overnight, and plated at 2.5 million cells/mL in RPMI medium in 6-well plates. Cells were stimulated with phorbol 12-myristate 13-acetate (PMA; 5 ng/mL, Sigma) and ionomycin (500 ng/mL, Sigma) or vehicle control (DMSO) for 6 h in the presence of monensin (2 µM, Sigma), brefeldin A (5 µg/mL, Sigma), and anti-CXCR3 antibody (BioLegend). After stimulation, cells were washed and stained with tetramers, viability dye, and surface antibody staining as above (Table S3). Intracellular staining for IL-2, TNF-α, IFN-γ, granzyme A, CD3, CD4, and CD8α was performed using the BD Cytofix/Cytoperm fixation/permeabilization kit according to the manufacturer’s instructions (BD). Cells were fixed in 2% paraformaldehyde, acquired on a spectral flow Cytek Aurora (ARC 1207i), and analyzed using FlowJo software (BD).

### Generation and analyses of T cell clones

#### Generation of T cell clones

Tetramer-positive cells were single-cell sorted into 96-well plates using purity mode on a FACSAria II (BD). Each well contained 10^5^ irradiated PBMCs and 10^4^ JY cells (Epstein–Barr virus– immortalized B lymphoblastoid cell line; Thermo Fisher Scientific), along with phytohemagglutinin (PHA; 1:100, Thermo Fisher Scientific), interleukin-7 (IL-7; 25 ng/mL, PeproTech), and IL-15 (25 ng/mL, PeproTech). IL-2 (50 IU/mL, PeproTech) was added on day 5 and replenished every 3–5 days. Cultures were restimulated every 2 weeks with fresh medium containing IL-2 (50 IU/mL), PHA (1:100), and 1 × 10^5^ irradiated PBMCs.

#### Generation of DCs

HLA-matched monocytes were differentiated into dendritic cells (DCs) in RPMI 1640 supplemented with glutamine, 10% FCS, penicillin–streptomycin, HEPES, GM-CSF (100 ng/mL), and IL-4 (500 U/mL). On day 3, fresh medium containing GM-CSF, IL-4, and 0.05 mM 2-mercaptoethanol was added. For peptide stimulation, immature DCs were harvested on day 5, plated at 0.1 to 0.25 million cells per well in flat-bottom 96-well plates, matured with LPS (100 ng/mL), and pulsed with peptides for 16–24 h prior to co-culture with T cells.

#### Stimulation of T cell clones

T cell clones were rested overnight in fresh media without IL-2, added to wells containing mature DCs, and incubated for 5 h in the presence of monensin (2 µM, Sigma-Aldrich) and brefeldin A (5 µg/mL, Sigma-Aldrich). Intracellular cytokine staining for TNF-α (BioLegend) was performed after stimulation using the BD Cytofix/Cytoperm Fixation/Permeabilization Kit according to the manufacturer’s instructions (BD).

### High-dimensional phenotypic analyses

SARS-CoV-2 tetramer–positive CD4^+^ T cells were manually gated, exported from FlowJo, and imported into R using the flowCore package. A total of 24 tetramer-positive populations were combined into a single dataset for downstream processing and high-dimensional analysis using the Spectre package in R ^29^. Marker intensities were transformed using an arcsinh transformation with a cofactor of 1900. Batch alignment was performed by initial coarse normalization using quantile-based scaling derived from a reference sample included in all batches, followed by application of the CytoNorm algorithm to adjust all samples ^43^. Clustering was performed using Phenograph with 30 nearest neighbors (k = 30). Uniform Manifold Approximation and Projection (UMAP) was used for dimensionality reduction and visualization^44^.

### Quantification and statistical analyses

Data transformation was performed using a base-10 logarithmic function for frequency, a base-2 logarithmic function for fold-change or inverse hyperbolic sine [sinh^−1^(X) = ln (X + sqrt (1 + sqr (X))) for MFI. Y=Bottom + X*(Top-Bottom)/(EC50 + X) was used to calculate EC50. Assessment of normality was performed using Shapiro-Wilk test. Spearman correlation was used if either of the two variables was nonnormal. Otherwise, Pearson correlation was used to measure the degree of association. The best-fitting line was calculated using simple linear regression. Statistical comparisons of two means were performed using paired or unpaired t test using a P value of < 0.05 as the significance level. Group comparisons were conducted using Fisher’s exact test, ANOVA, or mixed-effects models, as appropriate, with post hoc adjustment for multiple comparisons when overall tests were significant. Statistical analyses were performed using GraphPad Prism. Lines and bars represent mean, and variability is represented by SEM. Geometric means were computed as the exponential of the mean of log-transformed values. *P < 0.05, **P < 0.01, ***P < 0.001, ****P < 0.0001, and ns (not significant).

